# Pushing the limits of HiFi assemblies reveals centromere diversity between two *Arabidopsis thaliana* genomes

**DOI:** 10.1101/2022.02.15.480579

**Authors:** Fernando A. Rabanal, Maike Gräff, Christa Lanz, Katrin Fritschi, Victor Llaca, Michelle Lang, Pablo Carbonell-Bejerano, Ian Henderson, Detlef Weigel

## Abstract

Although long-read sequencing can often enable chromosome-level reconstruction of genomes, it is still unclear how one can routinely obtain gapless assemblies. In the model plant *Arabidopsis thaliana*, other than the reference accession Col-0, all other accessions *de novo* assembled with long-reads until now have used PacBio continuous long reads (CLR). Although these assemblies sometimes achieved chromosome-arm level contigs, they inevitably broke near the centromeres, excluding megabases of DNA from analysis in pan-genome projects. Since PacBio high-fidelity (HiFi) reads circumvent the high error rate of CLR technologies, albeit at the expense of read length, we compared a CLR assembly of accession Ey15-2 to HiFi assemblies of the same sample performed by five different assemblers starting from subsampled data sets, allowing us to evaluate the impact of coverage and read length. We found that centromeres and rDNA clusters are responsible for 71% of contig breaks in the CLR scaffolds, while relatively short stretches of GA/TC repeats are at the core of >85% of the unfilled gaps in our best HiFi assemblies. Since the HiFi technology consistently enabled us to reconstruct gapless centromeres and 5S rDNA clusters, we demonstrate the value of the approach by comparing these previously inaccessible regions of the genome between two *A. thaliana* accessions.

## INTRODUCTION

The first reference genome of *Arabidopsis thaliana*, from the accession Columbia (Col-0), was completed in the year 2000 with Sanger sequencing and assembled by a BAC minimal tiling path approach (1). Although it has served as the “gold standard” for the community ever since, it contains very little representation of the highly repetitive fraction of the genome, namely centromere repeats and ribosomal RNA genes. More than a decade later, the genomes of multiple other accessions were *de novo* assembled based on Illumina paired-end reads, but consisted of thousands of scaffolds (2–4). Notwithstanding their high error rate, long-read sequencing technologies, such as Oxford Nanopore Technologies (ONT) sequencing (reviewed in (5, 6)) and PacBio’s single-molecule real-time (SMRT) in the original continuous long read (CLR) sequencing mode (7), have significantly improved the contiguity of *de novo* assemblies. To date, there are 16 *A. thaliana* accessions sequenced with CLR technology (8–17), and although these assemblies commonly achieved some chromosome-arm level contigs, they invariably stop short of assembling through centromeric and pericentromeric regions. Only very recently, it was possible to assemble gapless centromeres in the *A. thaliana* reference accession Col-0, primarily with ultra-long ONT reads and the addition of PacBio high-fidelity (HiFi) reads for gap closing and polishing (18). Paradoxically, and despite rice (*Oryza sativa*) and maize (*Zea mays*) having complex genomes several times larger than *A. thaliana*, PacBio CLR technology has been successfully exploited to assemble gapless centromeres in about a third of the chromosomes in the pan-genome analyses of 31 rice (19) and 26 maize accessions (20, 21). This likely reflects fundamental differences in the composition of their centromeres. For instance, the tandem satellite repeats CentC (∼156 bp long) in maize are confined to a few small blocks interspersed with numerous centromeric retrotransposons (22). In contrast, the tandem *CEN180* satellite repeats (∼178 bp long) in *A. thaliana* Col-0 form very large arrays, only interrupted by 111 interspersed sequences larger than 1 kb (18).

PacBio HiFi reads, which are >99% accurate because they are generated from an innovative circular consensus sequencing strategy (23), overcome the high error limitation of ONT and CLR technologies at the cost of reducing read length. Recent studies in humans, rice and barley that compared HiFi-based assemblies to the other long-read technologies showed mostly an enhanced correctness, completeness and – sometimes – an improved contiguity (24–28). Those three metrics are often referred to as the “three C’s” and provide important information about the assembly quality. Among the most commonly used HiFi assemblers, both FALCON (11) and Canu (29) were originally conceived for PacBio CLR data. However, since the emergence of PacBio HiFi reads, FALCON added a HiFi-optimized parameter (23), while HiCanu emerged as a modification of the original Canu assembler (30). In contrast, Hifiasm (31), Peregrine (32) and IPA (33) were developed specifically for the purpose of assembling HiFi data.

Here, we compared genome assemblies resulting from a CLR library and the assemblies performed by five different state-of-the-art assemblers operating on a HiFi library of the same Arabidopsis accession, Ey15-2. We evaluated the impact of both coverage and read length in the metrics of contiguity, completeness and correctness, for which we analyzed a total of 255 HiFi assemblies based on subsets of the original HiFi data. We paid particular attention to the repetitive fraction of the genome and explored in detail the likely causes of contig breaks between both PacBio technologies and the different HiFi assemblers. Since the HiFi technology enabled us to obtain gapless centromeric regions, we present the first comparison of these previously unassembled regions of the genome between two *A. thaliana* accessions.

## RESULTS

To compare the performance of the current long-read sequencing platforms offered by PacBio, we generated CLR (subread coverage ∼1, 006x) and HiFi libraries (q20 HiFi read coverage ∼133x) starting from the same high-molecular-weight DNA extraction of a pool of individuals of the *Arabidopsis thaliana* natural accession Ey15-2 (accession ID 9994; CS76399) (Figure 1a). In addition, we produced an optical map with the Bionano Direct Label and Stain (DLS) technology (molecule coverage ∼781x) to validate and scaffold the main assemblies, and an Illumina PCR-free paired read library (coverage ∼166x) to evaluate completeness and accuracy of all assemblies, and to estimate the genome size of this particular strain.

**Figure 1.**
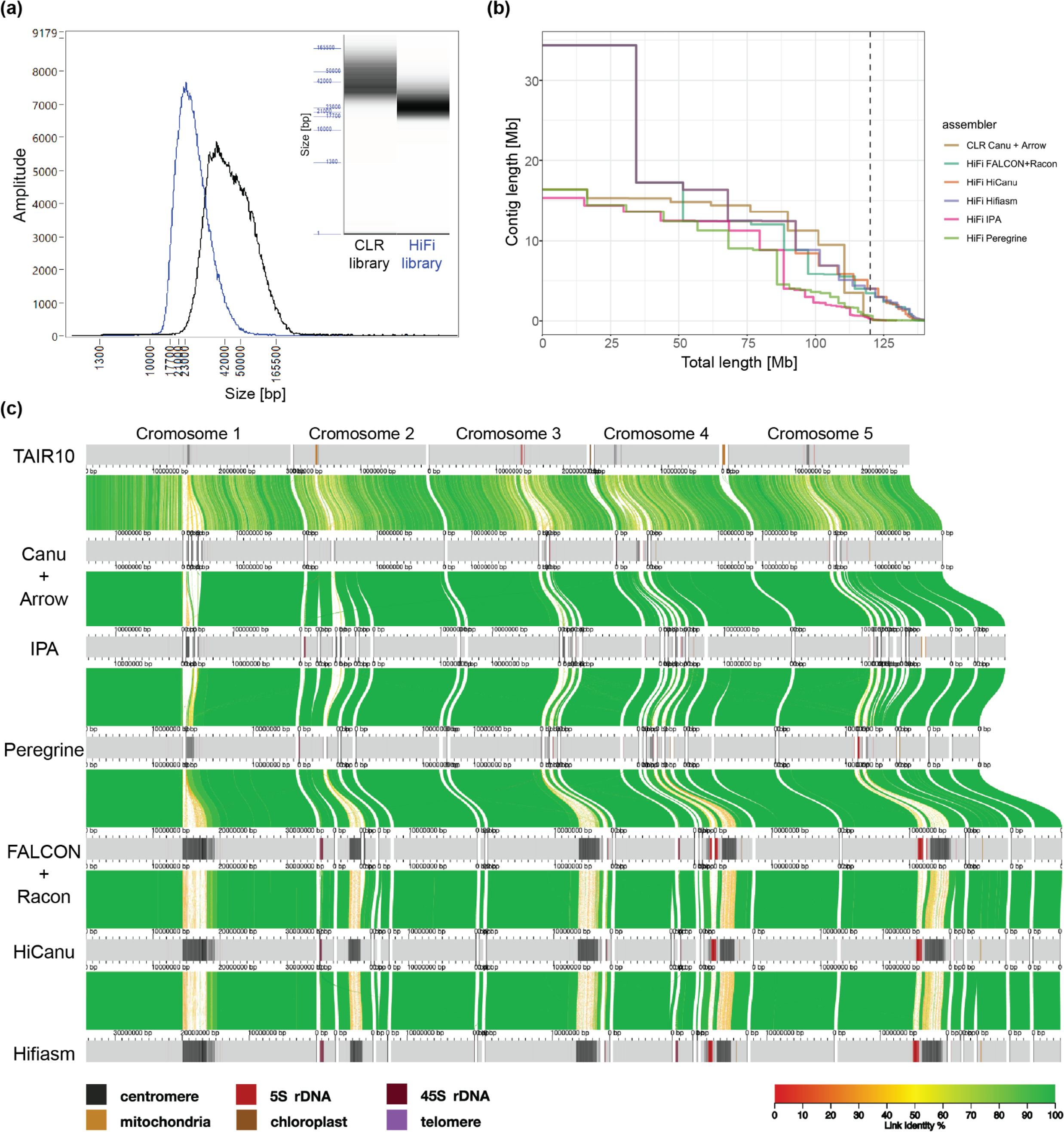
Comparison of different libraries and assemblers. **(a)** Insert size distribution of the CLR (black) and HiFi (blue) libraries after size-selection on the BluePippin instrument as measured on a Femto Pulse System. **(b)** Contiguity plot comparing the CLR and five HiFi assemblies using the complete dataset. For each assembly, the cumulative contig length (ordered from largest to shortest) is plotted over the estimated genome size of *A. thaliana* accession Ey15-2 (∼140 Mb). The vertical dashed line indicates the size of the TAIR10 reference genome. **(c)** Alignment of the TAIR10 reference genome and the contigs of the CLR and five HiFi assemblies visualized by AliTV (35). Co-linear horizontal gray bars represent chromosomes or contigs, with sequence annotated as repetitive elements (centromeres, 5S and 45S rDNAs, telomeres, mitochondrial and chloroplast nuclear insertions) displayed as shades. Only Bionano-scaffolded contigs >150 kb are shown. Distance between ticks equals 1 Mb. Colored ribbons connect corresponding regions in the alignment.

### Performance of the assembler of choice

To assemble contigs with the CLR dataset, we used Canu with a maximum input coverage of 200x, only using subreads larger than 10 kb, and polished the resulting assembly with Arrow (34), also using 200x of the initial long-reads. The resulting contigs had an NG50 of 14.82 Mb, which is on a par with the best published *Arabidopsis thaliana* CLR contigs (12–17).

With the HiFi dataset, we compared the performance of five different assemblers: FALCON (23), HiCanu (30), Hifiasm (31), Peregrine (32), and Pacbio’s Improved Phased Assembler (IPA; (33)). With the complete q20 HiFi dataset (∼133x), which has a median read length of 21.5 kb, we observed substantial differences in contig continuity for the different assemblers (Table 1). Only HiFi-Hifiasm and HiFi-HiCanu, both with 16.33 Mb, showed a higher NG50 than the CLR contigs. However, NG50 alone may not reflect the most noticeable differences in continuity between assemblers. HiFi-IPA and HiFi-Peregrine largest contigs, 15.33 Mb and 16.34 Mb, respectively, are comparable to the largest CLR-Canu contig (16.37 Mb), which represents an entire chromosome arm (Figure 1b). In contrast, HiFi-FALCON, HiFi-HiCanu and HiFi-Hifiasm all assembled a 34.36 Mb contig that corresponds to the telomere-to-telomere assembly of Chromosome 1 in *A. thaliana* (Figure 1c). The second largest contig was also exclusively assembled by those three assemblers. With 17.2 Mb, it spans the upper arm of Chromosome 3, presumably the entire centromere, and part of the other arm (Figure 1c). Similarly, the third largest contig of 16.33 Mb, only achieved by HiFi-Hifiasm and HiFi-HiCanu, corresponds to the upper arm of Chromosome 5, presumably encompassing the complete centromere, and part of the other arm (Figure 1c).

**Table 1.** Metrics of the CLR and five HiFi genome assemblies of A. thaliana Ey15-2.

| Assembler | Total length [Mb] | Scaffolded length* [Mb] | Largest contig [Mb] | Contig NG50 [Mb] | BUSCO completeness* | Merqury completeness* | QV* |
| --- | --- | --- | --- | --- | --- | --- | --- |
| CLR<br>Canu+<br>Arrow | 129.64 | 121.21 | 16.37 | 14.82 | C:98.7%<br>[S:98.0%,D:0.7%],<br>F:0.3%,M:1.0% | 98.72% | 54.45 |
| HiFi IPA | 125.96 | 123.43 | 15.33 | 12.41 | C:98.6%<br>[S:97.9%,D:0.7%],<br>F:0.3%,M:1.1% | 98.43% | 54.11 |
| HiFi<br>Peregrine | 295.26 | 124.98 | 16.34 | 11.28 | C:98.6%<br>[S:97.9%,D:0.7%],<br>F:0.3%,M:1.1% | 98.80% | 50.83 |
| HiFi<br>FALCON+<br>Racon | 140.60 | 136.09 | 34.35 | 12.44 | C:98.5%<br>[S:97.8%,D:0.7%],<br>F:0.4%,M:1.1% | 98.92% | 52.83 |
| HiFi<br>HiCanu | 234.77 | 135.57 | 34.36 | 16.33 | C:98.4%<br>[S:97.6%,D:0.8%],<br>F:0.4%,M:1.2% | 98.93% | 57.56 |
| HiFi<br>Hifiasm | 184.89 | 136.16 | 34.36 | 16.32 | C:98.5%<br>[S:97.7%,D:0.8%],<br>F:0.4%,M:1.1% | 98.94% | 60.26 |

We define NG50 as the sequence length of the shortest contig at 50% of the size of the TAIR10 reference genome (119.14 Mb; (38)). Benchmarking Universal Single-Copy Orthologs (BUSCO) (37) scores were obtained with the 'embryophyta\_odb10' set (n=1,375). Complete (C), single copy (S), duplicated (D), fragmented (F) and missing (M) genes are indicated. \*Scaffolded length, BUSCO scores, as well as Merqury's QV and completeness (36) were computed on contigs scaffolded with Bionano optical maps.

The total contig lengths of the different assemblers varied massively (Table 1), even among the HiFi methods, which have as input the exact same read set. Therefore, to evaluate accuracy and completeness on a more level playing field, we generated hybrid scaffolds of nuclear chromosomes for each of the described contig sets with Bionano optical maps. The scaffolded length of the different assemblers still differed by up to 14.95 Mb, equivalent to over 10% of the estimated genome size (see below), with the CLR-Canu, HiFi-IPA and HiFi-Peregrine assemblies at the low end, and the HiFi-HiCanu, HiFi-FALCON and HiFi-Hifiasm at the upper end (Table 1). By comparing k-mers in the *de novo* assemblies to those found in the raw PCR-free Illumina short reads, Merqury estimates base-level accuracy and completeness (36). The HiFi-Hifiasm assembly showed the highest accuracy, with a consensus quality (QV) score of 60.3, followed by HiCanu (QV 57.6). In contrast, the HiFi assemblers HiFi-IPA (QV 54.1), HiFi-Peregrine (QV 50.8) and HiFi-FALCON (QV 52.8) were all below the accuracy of the CLR-Canu assembly (QV 54.5). Meanwhile, k-mer based completeness was less informative due to little variation among assemblies, despite the massive variation in scaffolded length (Table 1). This is due to the fact that Merqury counts distinct k-mers found in the reads, regardless of their copy number (36). Similarly, the assessment of gene content of the assemblies with the widely used Benchmarking Universal Single-Copy Orthologs (BUSCO) score (37), although high (>98.4%), shows little difference among assemblies (Table 1). Therefore, practically all assemblers are successful in the non-repetitive fraction of the genome, but the repetitive regions are what deserve special consideration (see below).

### Impact of coverage

Since the HiFi technology enables the use of barcodes to sequence several samples per SMRTcell (39), it might in future become preferable to devote less read depth for *de novo* assembly applications. To simulate data sets with decreasing coverage, starting from our complete q20 HiFi dataset at 133x, we generated subsets – five replicates each – equivalent to 125x, 100x, 75x, 50x and 25x (Figure 2a). Each subset of reads was assembled with all five HiFi assemblers investigated in this study.

**Figure 2.**
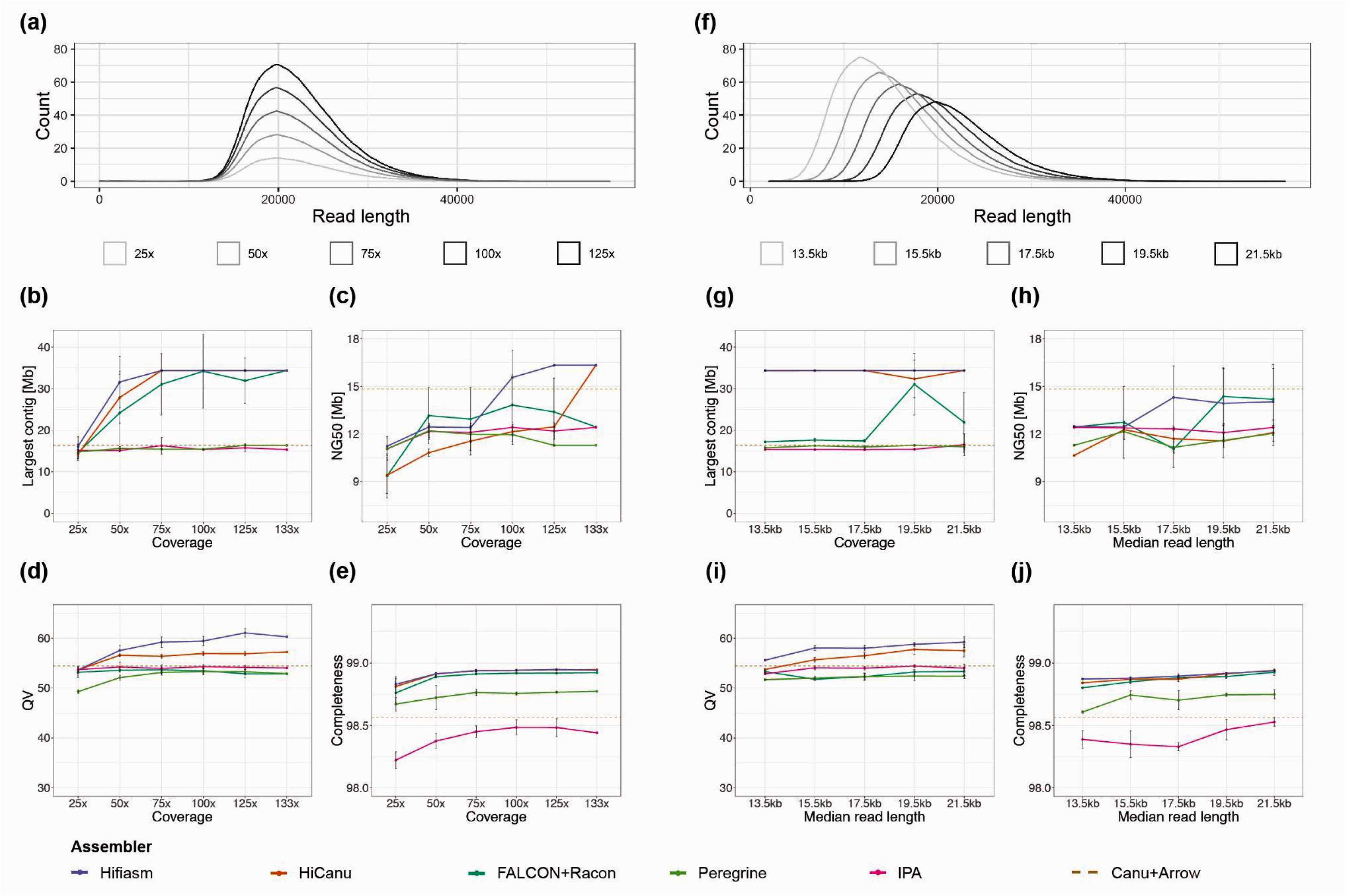
Impact of coverage and read length on assembly metrics. Read length distribution of subsets of HiFi reads with varying coverages: 25x, 50x, 75x, 100x and 125x **(a)**, and median read lengths: 13.5 kb, 15.5 kb, 17.5 kb, 19.5 kb and 21.5 kb **(f)**. Largest contig as a function of input coverage **(b)** and median read length **(g)**. Contig NG50 as a function of input coverage **(c)** and median read length **(h)**. We define NG50 as the sequence length of the shortest contig at 50% of the size of the TAIR10 reference genome (119.14 Mb; (38)). Consensus quality (QV) estimated by Merqury (36) as a function of input coverage **(d)** and median read length **(i)**. K-mer completeness estimated by Merqury (36) as a function of input coverage **(e)** and median read length **(j)**. QV and completeness were computed after reference-based scaffolding with RagTag (40).

Both HiFi-Hifiasm and HiFi-HiCanu successfully assembled the largest contig (∼34.4 Mb) in all replicates of subsets down to 75x coverage (Figure 2b). At 50x coverage, HiFi-Hifiasm failed to assemble this contig in one out of five replicate subsets, while with HiFi-HiCanu the contig broke in three of the replicates (Supplementary Figure 1). The lower continuity in HiFi-HiCanu when compared to HiFi-Hifiasm was also manifested in how often the second and third longest contigs were assembled, which is reflected by the progressive drop in NG50 at lower coverages (Figure 2c). Although HiFi-FALCON successfully assembled the three longest contigs in some replicates of subsets down to 50x coverage (Figure 2b), NG50 declined already at higher coverage than with HiFi-Hifiasm (Figure 2c). In addition, assemblies with HiFi-FALCON were more inconsistent across replicate subsets, to the degree that in two replicates of subset 100x chimeric contigs were formed (Supplementary Figure 2a-b). Nevertheless, HiFi-FALCON still performed better than both HiFi-Peregrine and HiFi-IPA in both continuity metrics. When compared to the CLR-Canu assembly, however, only HiFi-HiCanu with the full set and HiFi-Hifiasm with coverages at least 100x show a superior NG50 (Figure 2c).

After scaffolding, this time with RagTag (40), a reference-based scaffolding tool, we evaluated accuracy and completeness as described before. For all assemblers, QV scores were largely unaffected by coverage (Figure 2d), with HiFi-Hifiasm leading and HiFi-HiCanu coming in second. Only at 25x coverage, HiFi-Hifiasm and HiFi-HiCanu base-accuracy dropped to lower levels, but still comparable to all other assemblers, while HiFi-Peregrine QV scores fell below 50. Similarly, k-mer completeness was also largely unaffected by coverage with only slight drops at 25x for all HiFi assemblers (Figure 2e). Overall, HiFi-Hifiasm and HiFi-HiCanu stand out as the best assemblers across all metrics. In addition, HiFi-Hifiasm was more consistent in continuity and base quality, with compromises only apparent in some replicates of subsets with 75x and lower coverage.

### Impact of read length

Although the recommended insert size for HiFi libraries is 15-18 kb, we generated a q20 HiFi dataset with a median read length of 21.5 kb and N50 of 22.58 kb. This enabled us to simulate datasets – also five replicates each – with decreasing median insert sizes in steps of 2 kb down to 13.5 kb. With this, we could evaluate the impact of read length on various assembly metrics (Figure 2f). Due to the dependence on coverage observed before, all subsets were reduced to the highest common coverage (85x) to equalize the input conditions.

Both HiFi-Hifiasm and HiFi-HiCanu successfully assembled the largest contig representing Chromosome 1 in nearly all replicates of the different median read lengths, except for one instance, HiFi-HiCanu at 19.5 kb median read length (Figure 2g). HiFi-FALCON assembled the largest contig in half of the replicates of the two largest read length subsets, and failed to assemble it for all subsets with a median read length of 17.5 kb and shorter (Supplementary Figure 3). Similar to the situation observed in the coverage subsets, HiFi-FALCON produced a chimeric contig in one replicate of the subsets with median read length of 19.5 kb (Supplementary Figure 2c). The average NG50 produced by each HiFi assembler in all subsets was below the one achieved with CLRs (Figure 2h), which reflects the difficulty to assemble the second and third largest contigs (Supplementary Figure 3). HiFi-Hifiasm and HiFi-FALCON achieved higher average NG50 than the other HiFi assemblers for the two largest read length subsets, but NG50 dropped for HiFi-FALCON at 17.5 kb, and for HiFi-Hifiasm at 15.5 kb (Figure 2h). Both HiFi-Peregrine and HiFi-IPA did not show much variation either for the largest contig or NG50 across different read length subsets, and remain the HiFi assemblers performing the poorest for these metrics. Base-level accuracy and completeness for each assembler were very similar across all read length subsets, and the order mirrored what was observed for the complete read set (Figure 2i-j).

### Repetitive elements in scaffolds and contigs

To characterize the contribution of different genetic elements to the scaffolded genome for each of the assemblers, we annotated the repetitive elements in all contigs generated from the complete q20 HiFi dataset: transposable elements (TEs), centromeres, telomeres, 5S and 45S ribosomal RNA genes (rDNAs), as well as chloroplast and mitochondrial genome DNA insertions. In addition, through a k-mer based approach that employs Illumina PCR-free short reads (41), we estimated the nuclear genome size of this natural strain to be 143 Mb. Notably, the amount of non-repetitive sequence (understood as everything that was not annotated as a repetitive element) were very similar in the contigs successfully scaffolded with optical maps for the CLR and the HiFi assemblies (Figure 3a). While for the CLR the total non-repetitive sequence was 99.43 Mb (69.47%), for the HiFi assemblies it ranged from 98.99 Mb (69.16%) in HiFi-IPA to 100.47 Mb in HiFi-Hifiasm (70.20%). Even when adding telomeres, organellar insertions and TEs to the non-repetitive sequence, this length added up to only 118.97 Mb (83.12%) in the CLR-Canu assembly, while in the HiFi assemblies it ranged from 118.3 Mb in HiFi-IPA (82.65%) to 119.95 Mb (83.81%) in HiFi-Hifiasm (Figure 3a). These values are remarkably similar to the total length of 119.14 Mb of the TAIR10 reference genome (38).

**Figure 3.**
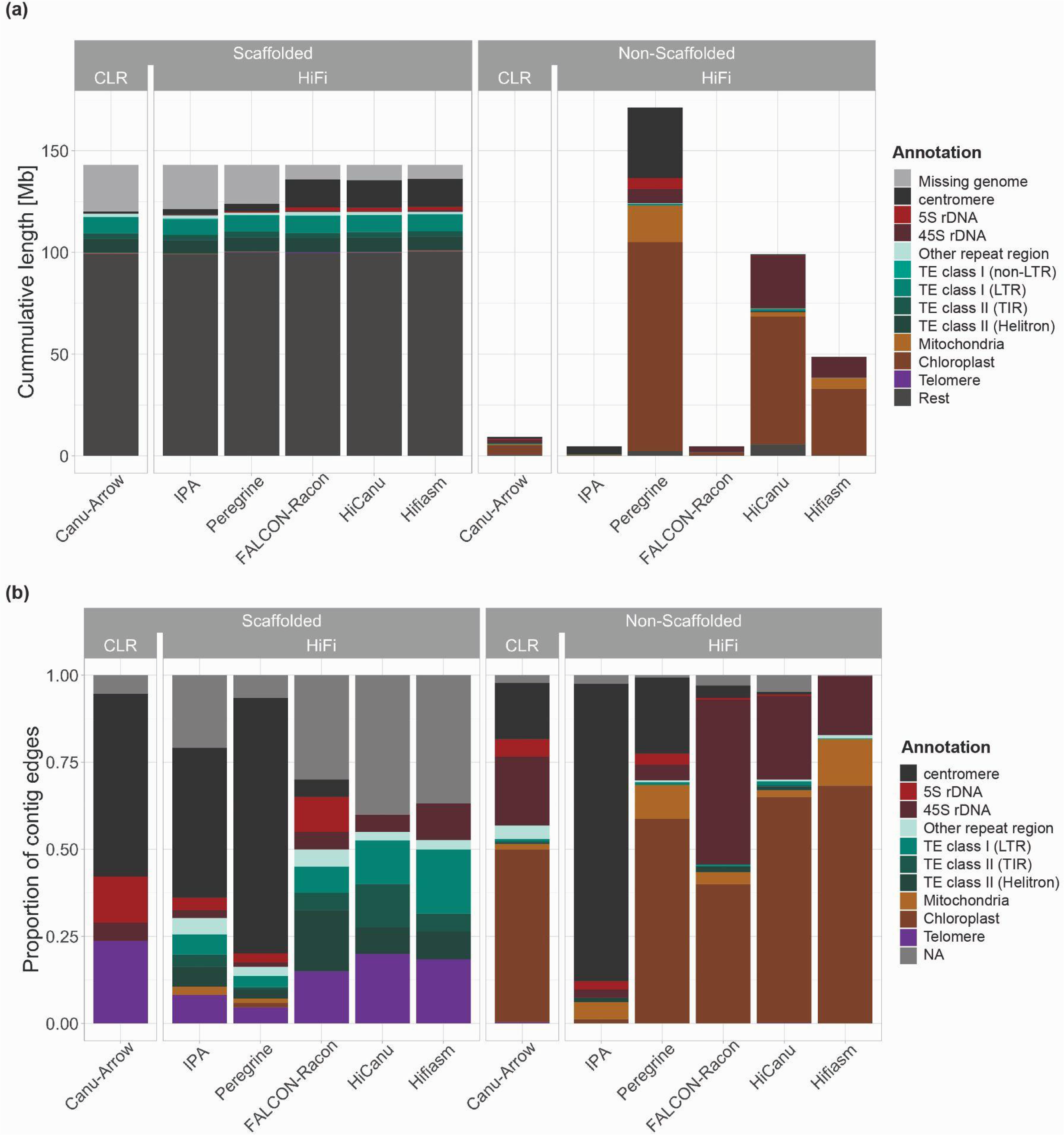
Repetitive elements in scaffolded and non-scaffolded contigs. **(a)** Stacked barplot of the cumulative length of various repetitive elements split into the scaffolded nuclear genome (left) and non-scaffolded contigs (right) for the CLR and HiFi assemblies. The height of the bars for the scaffolded genome is 143 Mb, the k-mer based genome size estimate by findGSE (41). **(b)** Fractions of the repetitive element found first within 2 kb of each contig edge in scaffolded contigs (left) and non-scaffolded contigs.

The substantial differences in the total length of nuclear scaffolds between technologies or assemblers are explained only when considering 5S rDNAs and centromeres. For the CLR-Canu assembly, we were only able to scaffold 159 kb of 5S rDNAs and 1.08 Mb of centromeres. Similar to the situation with other assembly metrics, performance of both HiFi-Peregrine and HiFi-IPA was closer to CLR-Canu than to the other HiFi assemblers. On the other hand, HiFi-FALCON, HiFi-HiCanu and HiFi-Hifiasm nuclear scaffolds contained 1.64-1.68 Mb of 5S rDNA and 13.63-13.69 Mb of centromeres. Therefore, the access to Mb-scale centromeric sequence and 5S rDNA arrays is what differentiates the most complete HiFi scaffolded assemblies from the CLR-based one (Figure 3a).

Nevertheless, even the largest scaffolded assemblies, i.e., Hifiasm, HiCanu and FALCON, do not reach the k-mer based genome size estimate; for these, there remain 6.94-7.52 Mb to be explained. To account for the missing sequence, we examined the non-scaffolded contigs. Their cumulative length per assembly (range 4.67 Mb to 171.29 Mb) varied much more dramatically than their scaffolded counterpart (Figure 3a). Most of these discrepancies can be attributed to the amount of organellar contigs. Similarly, the various assemblers produced discordant amounts of sequence annotated as 45S rDNAs, the length of which did not correspond to the difference between the genome size estimate and the lengths of scaffolded contigs for each assembly (Supplementary Figure 4). Notably, for the HiFi-Hifiasm assembly, with 10.36 Mb of non-scaffolded 45S rDNA, which represented 96% of the non-scaffolded sequence when removing organellar DNA, this value was different by only 3.42 Mb. To generate an independent 45S rDNA copy number estimate, we used a mapping-to-reference approach with Illumina PCR-free short reads (42), and estimated 1, 055 18S rRNA gene copies per haploid genome. Assuming 10.7 kb per 45S rDNA unit, this would equate to 11.28 Mb. Coincidentally, the amount of scaffolded and non-scaffolded 45S rDNA added up to 11.3 Mb. However, it is important to consider that since the non-scaffolded contigs consisting of 45S rDNA are not anchored to the assembled genome by non-repetitive sequence, it is very difficult to validate them. Unfortunately, when it comes to 45S rDNA clusters in *A. thaliana*, the high quality optical maps generated with the Bionano DLS technology are of limited use. This is due to the recognition sequence of the non-nicking enzyme DLE-1 (CTTAAG) occurring three times within 949 bp in the highly conserved 25S rRNA gene, while there are no occurrences in the more variable internal or external transcribed spacers of a reference 45S rDNA unit (43). This makes optical maps uninformative at these loci, in turn impeding the reliable construction of hybrid scaffolds.

### Where do contigs break?

To investigate in more detail the genetic elements that may cause contigs to break, we determined which of the annotated repetitive elements was found first within 2 kb of each contig edge. In an ideal case scenario, given that *A. thaliana* has five nuclear chromosomes, one would expect ten contig edges identified as telomeric repeats. In the CLR-Canu assembly, centromeric sequences were identified in more than half of the scaffolded contig edges (Figure 3b). Similarly, in the HiFi-Peregrine and HiFi-IPA assemblies, centromeric sequences at scaffolded contigs edges were found more often than any of the other repetitive elements (Figure 3b). In contrast, in the HiFi-FALCON assembly, only two scaffolded contig edges contained centromeric sequences while neither the HiFi-HiCanu nor the HiFi-Hifiasm contig breaks seemed to be due to centromeric sequence.

The next problematic repetitive elements for scaffolded contig edges in the CLR-Canu assembly were 5S rDNAs, followed by 45S rDNAs. At scaffolded contig edges, 5S rDNAs were also present in HiFi-IPA, HiFi-Peregrine and HiFi-FALCON assemblies, but not in HiFi-HiCanu and HiFi-Hifiasm. Regardless of the sequencing technology or assembler, all contigs that correspond to the upper arms of Chromosomes 2 and 4 broke at the subtelomeric 45S rDNA repeats (44). Contrary to the CLR assembly, all HiFi assemblies contain TEs in a substantial fraction of their scaffolded contigs edges (Figure 3b). We explain the underlying cause of these and most other contig breaks by analyzing more in detail the HiFi-Hifiasm assembly in the following section.

### In the quest of telomere-to-telomere assemblies

A major goal for *de novo* genome assembly projects is to achieve chromosome-level, telomere-to-telomere assemblies. Generally, the addition of orthogonal approaches (i.e., Hi-C, optical maps) is regarded as necessary to build confidence in the assembly (28). We compared whether this goal is within reach for our CLR assembly and our best HiFi (Hifiasm) assembly, when either is combined with optical maps.

The CLR-Canu assembly scaffolded with optical maps (without the aid of a reference-based scaffolding) does not achieve a single chromosome level assembly. Instead, ten hybrid scaffolds corresponding to complete chromosome arms, of which only three of them were slightly larger than the original contigs, plus two additional hybrid scaffolds confirming partial centromeres is the best outcome from these technologies combined (Figure 4a). In fact, only very seldom do Bionano DLS optical maps span complete *A. thaliana* centromeres (1001G+ Project, *personal communication*). For species for which there is a reference genome available, such as *A. thaliana*’s TAIR10, this limitation is not an issue, since reference-based scaffolding methods can be used to assign scaffolds to chromosomes. However, for species that lack a reference genome, Hi-C might be a better alternative to identify chromosome arms pairs.

**Figure 4.**
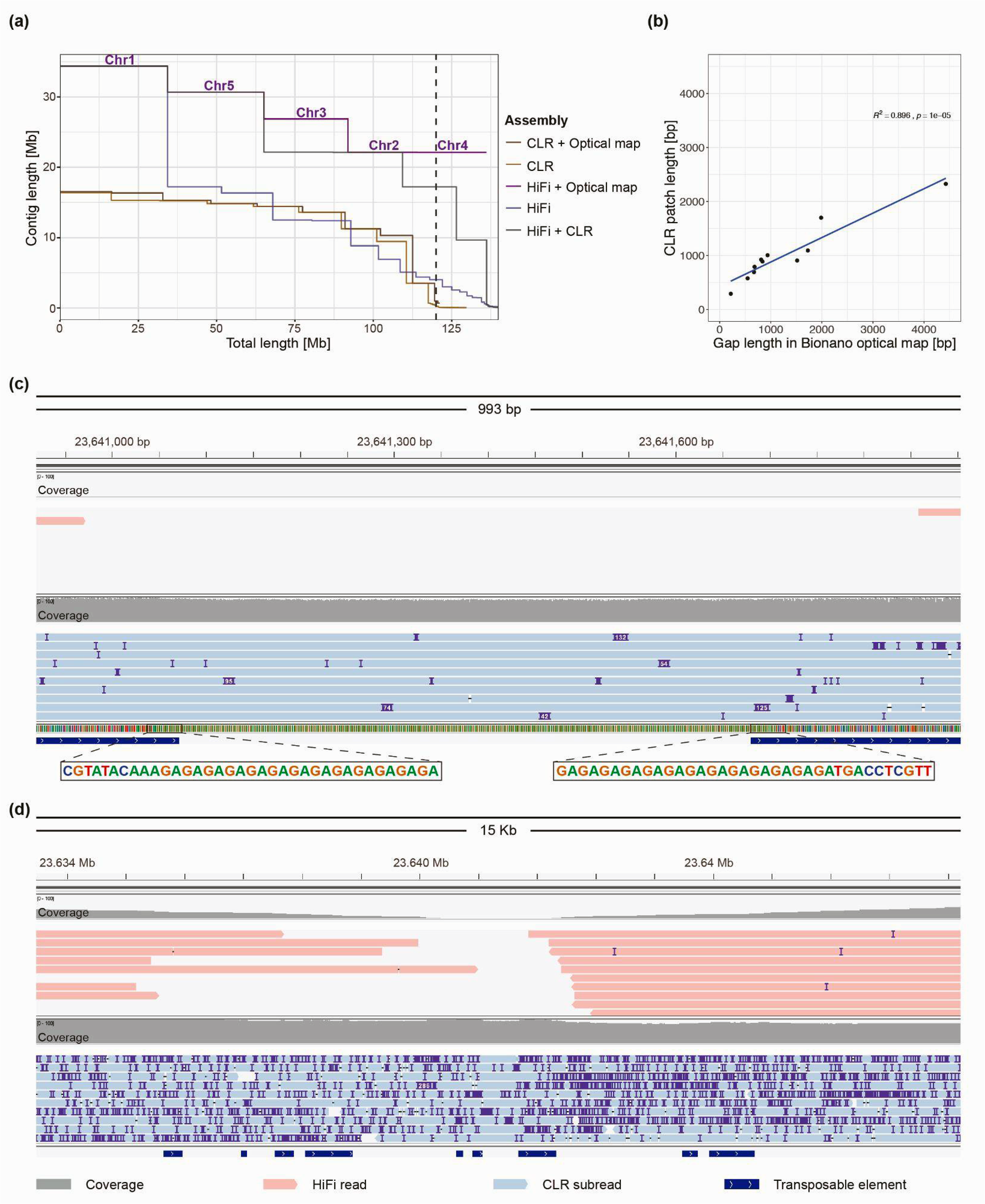
Hybrid assemblies and close inspection of gaps. **(a)** Contiguity plot comparing the CLR-Canu and HiFi-Hifiasm assemblies alone, combined with RagTag “patch” (40) or as hybrid scaffolds with Bionano optical maps. For each assembly, the cumulative contig – or scaffold – length (ordered from largest to shortest) is plotted over the estimated genome size of *A. thaliana* accession Ey15-2 (∼140 Mb). The vertical dashed line indicates the size of the TAIR10 reference genome. For the assembly that achieved “telomere”-to-telomere status (HiFi + Optical map), chromosome numbers are indicated on top of the scaffold lines. **(b)** Correlation of gap lengths estimates between Bionano optical maps and CLR “patches” introduced in the HiFi assembly by RagTag (40). **(c)** Visualization with IGV (46) of aligned HiFi reads (in red; top) and CLR (in blue; bottom) over Chr5:23640917-23641913, a locus in the HiFi + CLR assembly “patched” with the CLR-Canu assembly. **(d)** Zoom out of (c).

On the other hand, the HiFi-Hifiasm assembly combined with optical maps achieved five “telomere”-to-telomere hybrid scaffolds (Figure 4a), where the quotes in “telomere” indicates that the top of Chromosomes 2 and 4 ended after few dozens units of subtelomeric 45S rRNA genes. As shown in the analysis of contig breaks, all centromeres are complete in the HiFi-Hifiasm assembly (Figure 3b), while the remaining six fragmented chromosome arms (Figure 1c) were properly scaffolded, although with fourteen gaps. From these, twelve gaps had estimated sizes ranging from 217 to 6, 900 bp, and the other two were instead caused by contig overlaps not properly resolved by Hifiasm in Chromosomes 2 and 5. Contrary to the contig overlaps in Chromosome 5 (Supplementary Figure 5a), the optical map indicated that one of the contig edges in chromosome 2 was inconsistent for DLE-1 recognition sites (Supplementary Figure 5b). The conflicting contig edge contained two 45S rDNA units supported by a single – likely chimeric – HiFi read. Upon removal of this read and further re-assembly, the resulting scaffold contained a normal gap at this position.

Given that the CLR-Canu and the HiFi-Hifiasm contigs display mostly non-overlapping breaking patterns (Figure 1c), we combined both assemblies by preserving the most complete HiFi contig set and “patched” it with the CLR-Canu contigs using RagTag (40). This approach rendered four “telomere”-to-telomere chromosomes, with Chromosome 3 split into two scaffolds (Figure 4a) separated by a gap estimated to be 6, 900 bp according to the optical map (Supplementary Figure 5c). The pair of overlapping HiFi contigs belonging to Chromosome 5 was also identified and corrected by RagTag, which removed 7 bp (Supplementary Figure 5d). The CLR assembly only contributed a total of 12, 049 bp distributed in twelve “patches”, ranging from 290 to 2, 326 bp, largely in agreement with the gap sizes previously estimated by the optical map (Figure 4b; Supplementary File 1). A closer examination into these “patches” revealed that all consisted of either GA/TC or GAA/TTC low-complexity repeats as evidenced by the subreads of the CLR library that spanned the complete region without a noticeable drop in coverage (Figure 4c). In contrast, q20 HiFi reads showed a drop in coverage extending for several kilobases around the low-complexity repeats (Figure 4d), which were generally not covered by any read – or by a single read in three out of the twelve instances.

This coverage bias of the HiFi chemistry at GA/TC low-complexity repeats was previously noticed for four out of twelve gaps of a Chromosome X in humans (30). To investigate whether this particular class of low-complexity repeats is responsible for contig breaks in a different *A. thaliana* genome, we re-sequenced with HiFi reads and assembled with Hifiasm a single individual of the accession Col-0 (accession ID 6909; CS76778). The reference-based scaffolds contained only nine gaps. A comparison of our HiFi Col-0 assembly with the TAIR10 reference genome (38) and two recently published Col-0 (18, 45) assemblies confirmed that eight of the nine gaps (range: 601 to 1, 861 sizes) also occurred at GA/TC or GAA/TTC repeats (Supplementary File 1), with the remaining gap consisting of an unresolved 42, 895 bp overlap between two contigs – when compared to the Naish *et al.* assembly (18). That contigs breaks in these *A. thaliana* assemblies were mostly due to GA/TC low-complexity repeats (85.71% and 88.88% in the Ey15-2 and the Col-0 assemblies, respectively) denotes a current limitation of HiFi reads, however, the relatively small sizes of the gaps they incurred is rather encouraging for the technology.

### Natural variation in centromeres and 5S rDNA clusters

Two recently published assemblies of the reference accession Col-0 have fully (18) or partially (45) resolved centromeres. Since our HiFi assemblies also provide access to previously unassembled regions of the nuclear genome (Figure 1c, Figure 3b), most notably, centromeres, 5S rDNA clusters and large insertions of organellar DNA, we compared these repetitive regions in our assembly of Ey15-2 with all existing assemblies of Col-0 (Figure 5a-b). Among the available Col-0 assemblies, there was high consistency in the length, orientation and overall structure for centromeres in Chromosomes 1, 3, 4 and 5 (Figure 5a; Supplementary Figure 6). Only the centromere of Chromosome 2 in the assembly from Wang *et al.* is slightly shorter, which could potentially be attributed to a gap in this assembly within the centromere (45).

**Figure 5.**
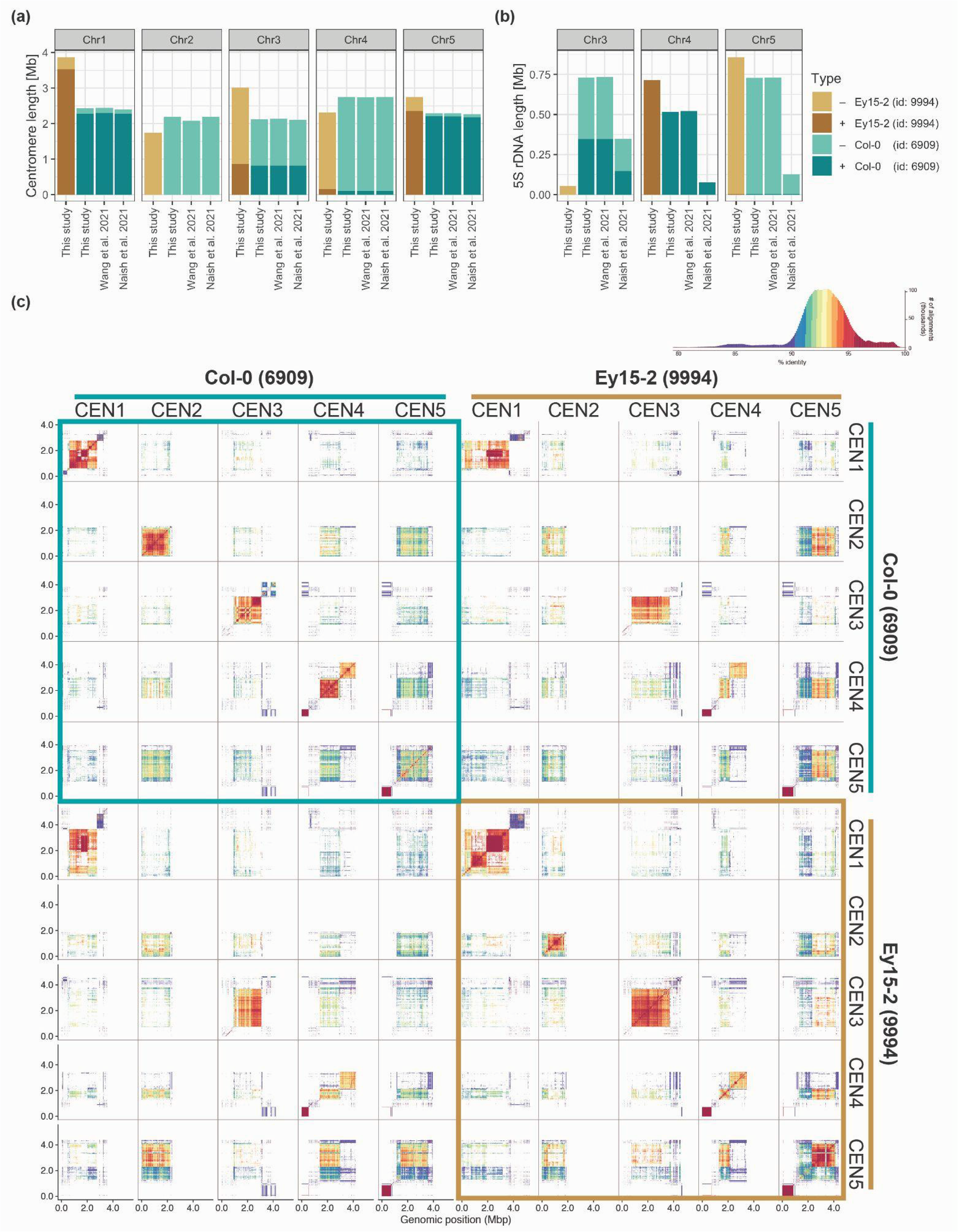
Centromere and 5S rDNA variation between *A. thaliana* accessions Ey15-2 and Col-0. **(a)** Centromere and **(b)** 5S rDNA length of each chromosome in the HiFi-Hifiasm assembly of accession Ey15-2 and three independent assemblies of accession Col-0: HiFi-Hifiasm in this study, ONT+HiFi in Wang *et al.* (45) and ONT+HiFi in Naish *et al.* (18). **(c)** Comparison of all pericentromeric regions in the HiFi-Hifiasm assemblies of Col-0 and Ey15-2 visualized by StainedGlass (47). A histogram of the colored percent identity is shown at the top-right of the panel.

In Col-0, CEN1 is the most different among the five centromeres (18, 45). Comparing our two accessions, CEN1 in Ey15-2 is at least 1.4 Mb longer than CEN1 in Col-0 (Figure 5a). Despite the length difference, CEN1 in Ey15-2 is more related to CEN1 in Col-0 than to any other Ey15-2 centromere (Figure 5c; Supplementary Figure 7a). In Ey15-2, there are two arrays encompassing CEN1, both larger than their counterpart in Col-0. The main array (upstream) consists of two distinct subarrays divided by a short inverted region (Supplementary Figure 7), and the downstream array is even more dissimilar to the other centromeres than the upstream one (Figure 5c). CEN2 is similar in size and orientation in Ey15-2 and Col-0, the latter being ∼450 kb larger (Figure 5a). CEN3 in Ey15-2 is ∼900 kb larger than in Col-0, the second largest size difference between homologous centromeres (Figure 5a). In spite of that, CEN3 of both accessions have the same inverted structure and they are also similar at the sequence level (Figure 5c). CEN4 is ∼440 kb larger in Col-0 than in Ey15-2 (Figure 5a). Like Col-0, Ey15-2 has a dual CEN4, with each array being very distinct to the other (Figure 5c). As in Col-0, the upstream array is more similar to the other chromosomes. The downstream array, however, shows more similarity to its counterpart in Col-0 than to any other centromere within Ey15-2 (Supplementary Figure 7b). Finally, CEN5 is > 460 kb longer in Col-0 than in Ey15-2 (Figure 5a). In the latter, there are various inversions throughout the entire array (Supplementary Figure 7).

Regarding the 5S rDNA clusters, while their size and orientation were highly consistent between our Col-0 HiFi assembly and the one from Wang *et al.* for Chromosomes 3, 4 and 5 (45), they were substantially smaller in the assembly from Naish *et al.* for all three loci (18) (Figure 5b). An important distinction between the previously published Col-0 genomes is that despite both being hybrid assemblies of ONT and Pacbio HiFi reads, the underlying contigs in Naish *et al.* are primarily ONT-based (18) while in Wang *et al.* they are ultimately HiFi-based (45). In a previous study, we have estimated the 5S rRNA gene copy number in Col-0 to be >2, 000 by quantitative PCR, and considered this an underestimate given that the primers may have missed units due to polymorphisms (48). With 1.98 Mb annotated as 5S rDNA, and considering that each 5S rDNA unit is ∼500 bp, our Col-0 HiFi assembly contains ∼3, 962 5S rRNA genes while that of Naish *et al.* only ∼1, 111. Since the Col-0 individual we sequenced originated from the exact same seed batch as those used by Naish *et al.* (18), and since 5S rRNA gene copy number has been shown to be rather stable in *A. thaliana* mutation accumulation lines propagated by single-seed descent (48), we speculate that this discrepancy likely reflects differences in the underlying long-read sequencing technologies (namely, PacBio HiFi versus ONT) and assembly algorithms, as opposed to a real biological difference between samples. To obtain a copy number estimate before the assembly process, we identified 5S rRNA genes directly on the q20 HiFi reads and, after normalizing by genome-wide read-depth, the estimate was 2, 983 copies. This is ∼1, 000 copies less than in the Col-0 HiFi assembly, but nearly 1, 900 more than in the assembly from Naish *et al.* While it remains challenging to determine the exact 5S rRNA gene number in the Col-0 genome, the latter estimate from unassembled long-reads is closer to both HiFi-based assemblies than to the ONT-based assembly.

When comparing the two different accessions, the orientation and size of the major 5S rDNA clusters in the upper arms of Chromosomes 4 and 5 are similar, and only slightly larger in Ey15-2 (Figure 5b). Also, the minor 5S rDNA cluster in the lower arm of Chromosome 5 is conserved (Supplementary Figure 8). In contrast, 5S rDNA repetitive elements only total 55 kb of Chromosome 3 in Ey15-2, that is, depending on whether we compared with the ONT-based or HiFi-based assemblies, six to thirteen times less than in Col-0. Presence/absence variation of 5S rDNA clusters in Chromosome 3 between *A. thaliana* accessions is well known from cytological studies (48–50). With telomere-to-telomere assemblies that fully resolve centromeric and pericentromeric regions, we can now add several layers of resolution to these comparisons. Besides characterizing the actual length and orientation of the polymorphic 5S rDNA clusters themselves (Figure 5b), we can better appreciate their genomic neighborhood. For instance, from the two 5S rDNA clusters in the lower arm of Chromosome 3 in Col-0 that are in different strand orientation, Ey15-2 only carries a minor version of the downstream cluster in the negative strand (Supplementary Figure 8).

As for organellar DNA insertions into the nuclear genome, the large mitochondrial DNA insertion downstream of the centromere in Chromosome 2 in Col-0 is absent in Ey15-2 (Supplementary Figure 8). Although this insertion remains only partially characterized in the TAIR10 reference genome, fiber-fluorescence *in situ* hybridization analyses have shown it is ∼620 kb long (51). The large mitochondrial DNA insertion represents another locus inconsistent among the three Col-0 assemblies. While in the assembly from Naish *et al.* (18) the size of the locus is 369 kb, in our HiFi-Hifiasm Col-0 assembly and the one from Wang *et al.* (*45*) it is 640 kb (Supplementary Figure 9), in remarkable agreement with the previous cytological estimate (51).

## DISCUSSION

Here, we have compared a CLR genome assembly that rivals the best published *A. thaliana* CLR assemblies with different HiFi assemblies produced with five state-of-the-art HiFi assemblers of the same sample. We find that a high-quality HiFi data set is preferable and, although a hybrid assembly of these two technologies accomplished a “telomere”-to-telomere genome with a single gap, only minor gains can be achieved by adding CLR data. An important insight is how much the choice of HiFi assemblers matters, to which we can confidently speak because we systematically compared their performance with the same long-read datasets. In our model organism, the HiFi assemblers FALCON, HiCanu and Hifiasm allowed us to access nearly 15 Mb more nuclear DNA sequence than the CLR assembly, primarily in the form of centromeres and 5S rDNA clusters (Figure 3a), with negligible differences in the non-repetitive fraction of the genome (Table 1). Hifiasm was our preferred choice because it achieved not only the highest consensus quality, but also because contiguity of the assembly was highly robust to a decrease in coverage and median read length (Figure 2).

Despite HiFi long reads supporting the successful assembly of centromeric regions, the contig breaks along several chromosome arms – thought to be less challenging than highly repetitive centromeres – were initially puzzling (Figure 1c). Many contigs that did not end with telomeres or 45S rDNA repeats, carried TEs at their edges, and several could at first not be explained (Figure 3b). PacBio CLR and ONT assemblies for the two HiFi genomes sequenced in this study helped us to shed light on the underlying cause for the vast majority of these breaks: GA/TC low-complexity repeats, which are poorly represented in the source HiFi reads (Figure 4c-d). Fortunately for future HiFi assemblies of *A. thaliana* genomes, the confirmed sizes of gaps due to this class of repeats were relatively small, ranging from 290 to 2, 326 bp (Figure 4b). We therefore strongly favor the HiFi technology for routinely obtaining chromosome-level assemblies with gapless centromeres without the need of complementary chromosome scaffolding techniques such as optical or chromosome contacts maps.

Based on the success of centromere assemblies, we are excited by the prospect of analyzing centromeres and 5S rDNA clusters from multiple accessions, given the intriguing observations we have already made in a comparison between Ey15-2 and Col-0. For example, it will be of interest to learn whether relatively conserved structural features, such as the bimodal centromere array in Chromosome 4, is common, or whether the downstream array, which presents low CENH3 occupancy in Col-0 (18), has diverged and been lost in other accessions. Similarly, it will be interesting to learn whether CEN1 stands apart in other accessions as well, or whether certain centromeres are more restricted in length variation. As for the 5S rDNA clusters, the full reconstruction of these loci in other accessions will enable the identification of cluster-specific polymorphisms that can serve as reporters of the expression status of each cluster, which could have implications on the 3D organization of chromatin within the nucleus.

## DATA AVAILABILITY

Raw data for the genome assemblies of *Arabidopsis thaliana* accessions Ey15-2 and Col-0, such as PacBio CLR and HiFi reads and Illumina PCR-free paired-end reads can be accessed in the European Nucleotide Archive (ENA; https://www.ebi.ac.uk/ena/browser/home) under the project accession number PRJEB50694. Custom scripts and small files to reproduce the analyses in this study can be found in the dedicated GitHub repository (https://github.com/frabanal/A.thaliana_CLR_vs_HiFi). Larger files, such as the hard-masked version of TAIR10, the main genome assemblies, annotation files and Bionano optical maps can be found on: https://keeper.mpdl.mpg.de/d/216caab287514b1ba2c5/.

## ACKNOWLEDGEMENT

We thank Haim Ashkenazy for helpful advice on assembly issues, Corinna Kersten for assistance in the visualization of optical maps, and members of the *Arabidopsis thaliana* 1001G+ Consortium for fruitful discussions.

## FUNDING

This work has been supported by a Human Frontiers Science Program (HFSP) Long-Term Fellowship (LT000819/2018-L) to FAR, by DFG-funded ERA-CAPS 1001G+, and by the Max Planck Society.

## CONFLICT OF INTEREST

The authors declare no competing or financial interests.

## AUTHOR CONTRIBUTIONS

Conceptualization, F.A.R.; Methodology, F.A.R., M.G., P.C.B.; Investigation, F.A.R., C.L., K.F., V.L., M.L.; Formal Analysis, M.G., F.A.R., V.L., I.H.; Resources, D.W., I.H.; Writing – Original Draft, M.G., F.A.R.; Writing – Review & Editing Preparation, F.A.R., D.W.; Visualization, F.A.R., M.G., I.H.; Supervision, F.A.R., D.W.; Project Administration, F.A.R.; Funding Acquisition, D.W., F.A.R.

## MATERIAL AND METHODS

### Plant growth conditions

*A. thaliana* seeds of the natural strains Ey15-2 (accession ID 9994; CS76399) and Col-0 (accession ID 6909; CS76778) were germinated on soil and stratified in darkness at 4°C for six days, after which they were transferred to long day conditions (16 h light) at 23°C and 65% relative humidity under 110–140 μmol m^-2^ s^-1^ light provided by Philips GreenPower TLED modules (Philips Lighting GmbH, Hamburg, Germany). To reduce starch accumulation, 21-days-old and 26-days-old plants of Ey15-2 and Col-0, respectively, were placed into darkness for 24 h before harvesting. For Ey15-2, ca. 30 g of flash-frozen rosettes from multiple individuals were ground in liquid nitrogen with pestle and mortar. For Col-0, a single individual was harvested and processed in a similar manner.

### High molecular weight DNA extraction

For Ey15-2, we extracted high molecular weight DNA (HMW-DNA) as described previously (16). Briefly, tissue powder was resuspended in 500 ml of freshly prepared and ice-cold nuclei isolation buffer (NIB: 10 mM Tris pH8, 100 mM KCl, 10 mM EDTA pH8, 500 mM sucrose, 4 mM spermidine, 1 mM spermine). The homogenate was filtered through two layers of miracloth (EMD Millipore; 475855-1R) and distributed in several 50 ml FALCON tubes, to which 1:20 (v/v) of NIB containing 20% Triton-X-100 was added. Samples were incubated on ice for 15 min., and centrifuged at 3, 000 g at 4°C for 15 min. Nuclei pellets were pooled together, washed with ca. 35 ml of NIB containing 1% Triton-X-100, and further centrifuged at 3, 000 g at 4°C for 15 min. The resulting pellet was gently resuspended in 20 ml of pre-warmed (37°C) G2 lysis buffer (Qiagen; Cat. no. 1014636), incubated with 50 μg/ml RNaseA (Qiagen; Cat. no. 19101) at 37°C for 30 min, followed by 200 μg/ml proteinase K treatment (Qiagen; Cat. no. 19133) at 50°C for 3 h. After centrifugation at 8, 000 g at 4°C for 15 min, the supernatant containing the DNA was purified with Genomic-tip 100/G (Qiagen; Cat. no. 10243) with the Genomic DNA Buffer Set (Qiagen; Cat. no. 19060) following the manufacturer’s instructions. To the resulting flow-through, 0.7 volumes of isopropanol were gently added, and the precipitated DNA was spooled with a glass hook through slow tube rotations, and resuspended in EB buffer (Qiagen; Cat. no. 19086) overnight at 4°C.

For Col-0, we extracted HMW-DNA following a modified version of the Mayjonade *et al.* (2016) protocol (52) that included the addition of ß-mercapto-ethanol during the lysis step and a Phenol:Chloroform purification step (53). Briefly, 300 mg of tissue powder was incubated for 45 min. at 55°C in freshly prepared and pre-heated lysis buffer (1% sodium metabisulfite, 1% PVP40, 0.5 M NaCl, 100 mM Tris HCl pH8, 50 mM EDTA pH8, 1.5% SDS, 2% ß-mercapto-ethanol). The following steps were performed at room temperature. 60 µL of 20 mg/ml PureLink^TM^ RNAseA (Thermo Fisher Scientific; Cat. No. 12091021) was added to the lysate and incubated for 10 min. To precipitate proteins, 600 µL of 5 M potassium acetate was added to the samples followed by 2.4 ml of 25:24:1 (v/v/v) Phenol:Chloroform:Isoamyl alcohol (ROTI^®^; Cat. No. A156.1) and incubated for 10 min. on a rotor. After centrifuging at 4, 400 x g for 10 minutes, the upper phase was transferred to a new tube and mixed with 24:1 (v/v) Chloroform:Isoamyl alcohol for 10 min. on a rotor. Following a second centrifugation step at 4, 400 x g for 10 min., the upper phase was transferred to a new tube and two bead cleanups were performed to remove contaminants. The first bead cleanup was performed for 30-60 min. with 1.0x volume of 0.4% solution of SeraMag SpeedBeads® Carboxyl Magnetic Beads (GE Healthcare), followed by an incubation of 30-60 min. on a rotor. After placing the tube on a magnet, the supernatant was discarded and beads were washed two times with 80% ethanol. Elution was performed with 50 µL EB (QIAGEN) after an incubation at 37 °C for 15 min. The second cleanup was performed with 0.45x volume of AMPure PB magnetic beads (P/N 100-265-900, Pacific Biosciences, CA). After a binding time of 30 min. on a rotator, beads were placed on a magnet and washed two times with 80% ethanol. For elution, 45 µL EB (QIAGEN) was added and incubated for 10-15 minutes on a rotor.

### Long-reads libraries preparation

For the CLR library of Ey15-2, 10 μg of 2x needle-sheared (FINE-JECT 26Gx1” 0.45×25 mm, LOT 14-13651) HMW-DNA was used for a double library prepared with the SMRTbell Express Template Preparation Kit 2.0 (P/N 101-693-800 Version 01, Pacific Biosciences, CA), and size-selected with the BluePippin system (SageScience) with 30 kb cutoff in a 0.75% DF Marker U1 high-pass 30-40kb vs3 gel cassette (BLF7510, Biozym). The library was sequenced in a single SMRT Cell (30 hours movie time) with the Sequel II system (Pacific Biosciences, CA) using the Binding Kit 2.0 (P/N 101-842-900).

For the HiFi library of Ey15-2, HMW-DNA (25 ng/μl) was separately sheared with 30 kb and 35 kb settings using a Megaruptor 2 instrument (Diagenode SA). However, the resulting average insert sizes were shorter than expected, approximately 19 kb and 24 kb, respectively. Therefore, 10 μg of both sheared fractions were combined in equal amounts and used for a double library (Procedure & Checklist: P/N 101-853-100 Version 03, Pacific Biosciences, CA) with the HiFi SMRTbell® Express Template Prep Kit 2.0, and size-selected with the BluePippin system (SageScience) with 17 kb cutoff in a 0.75% DF Marker S1 High-Pass 6-10kb vs3 gel cassette (BLF7510, Biozym). The library was sequenced in a single SMRT Cell (30 hours movie time) with the Sequel II system (Pacific Biosciences, CA) using the Binding Kit 2.0 (P/N 101-842-900).

For the HiFi library of Col-0, HMW-DNA (120 ng/µl) was sheared two times (back and forth) with a gTUBE (Covaris; P/N 520079) in a Eppendorf™ Centrifuge 5424 at 4800 rpm (soft) for 3 x 1 min. 5 µg of sheared DNA were used to prepare libraries using the HiFi SMRTbell® Express Template Prep Kit 2.0 (PN100-938-900) with SMRTbell Barcoded Adapter bc1022 (’CACTCACGTGTGATAT’) and SMRTbell Enzyme Clean Up Kit 2.0 (PN 101-932-600). Since this library was multiplexed with another unpublished sample, we used the protocol “Procedure & Checklist” (P/N 101-853-100 Version 04, Pacific Biosciences, CA) with some modifications. The two libraries were combined in equal amounts and size-selected with the BluePippin system (SageScience) with 10 kb cutoff in a 0.75% DF Marker S1 High-Pass 6-10kb vs3 gel cassette (BLF7510, Biozym). The library pool was sequenced with sequencing primer v5 (P/N 102-067-400) in a single SMRT Cell following the loading and pre-extension recommendations (P/N 101-769-100 version 6 to 9) with the Sequel II system (Pacific Biosciences, CA) using the Binding Kit 2.2 (P/N 101-894-200).

### DNA extraction and short-reads library preparation

DNA for PCR-free data was extracted with the DNeasy Plant Mini Kit (Qiagen; Cat. no. 69104) following the manufacturer’s instructions from the same ground tissue as the HMW-DNA. 700 ng of DNA were fragmented using a Covaris S2 Focused Ultrasonicator (Covaris) with settings: intensity 5, 10% duty cycle, 200 cycles and 45s treatment time. Subsequent library preparation was performed with the NxSeq® AmpFREE Low DNA Library Kit (Lucigen®; Cat. no. 14000-1) according to the manufacturer’s instructions with one slight modification. Following adapter ligation and prior to the final bead-cleanup at the purification step, we introduced an additional bead-cleanup (0.6:1, bead:library ratio) that serves as a large-cutoff to remove long inserts. Library concentration was measured with the Qubit® 2.0 Fluorometer (Invitrogen), and the insert size distribution was estimated to be around 460 bp (including adaptors) with a High Sensitivity DNA Chip (Agilent; Cat. no. 5067-4626) on an Agilent Bioanalyzer 2100 instrument. The library was sequenced as paired-end 150 bp reads to a coverage depth of ca. 166x on an HiSeq 3000 instrument (Illumina).

### Generation of optical map

*A. thaliana* plants of accession Ey15-2 were germinated *in vitro* and transferred to soil in flats. To minimize starch accumulation, plants were placed in the dark for 24 hours before tissue collection. Ultra-HMW DNA was isolated from young plants using a modified version of the protocol described in Deschamps *et al.* (2018) (54), which is based on the Bionano DNA Plant Isolation kit (Cat 80003; Bionano Genomics, San Diego, CA). Approximately 2 grams of young, healthy, light-starved leaves were transferred to a 50 ml conical tube and incubated for 20 min. in 60 ml ice-cold Bionano Fixing solution with added 3.2 ml formaldehyde, followed by three 10 min. washes in 60 ml ice-cold Bionano fixing solution without formaldehyde. The resulting fixed tissue was placed in a chilled square petri dish with 4.5 ml ice-cold Bionano Homogenization buffer supplemented with 1 μM spermine tetrahydrochloride, 1 μM spermidine trihydrochloride and 0.2% ß-mercapto-ethanol. The leaves were manually chopped with a razor blade and transferred to a 50 ml conical tube, blended 3 to 4 times for 20 sec. in ice using a Qiagen Tissue ruptor and filtered through 100 μM and 40 μM cell strainers. Nuclei and cell debris were pelleted by centrifugation at 3, 100 g, the supernatant decanted and the resulting pellet resuspended by swirling. Excess starch and cell debris in the original pellet were removed by low-speed centrifugation. The tube with the resuspended pellet was filled with fresh homogenization buffer, mixed by inversion and centrifuged for 2 min. at 100 g with slow deceleration. The top 75% of the supernatant was recovered by carefully decanting 35 ml into a new 50-ml tube, leaving excess contaminants at the bottom in the last 10-15 ml. This process was repeated 2 or 3 times until the supernatant was clear and the pellet had a reduced size. The nuclei in the supernatant were recovered by centrifugation at 3, 100 g and were resuspended in 55 μl cold Bionano Density Gradient Buffer. The tube containing the final resuspension was incubated at 43°C, mixed with 1X melted low-melting-point agarose equilibrated at 43°C and allowed to solidify after transferring to a plug mold. The agarose-embedded nuclei were incubated twice at 50°C in Bionano Lysis Buffer with added 8% (v/v) Puregene proteinase K, for a total of 12-16 h. Puregene RNase A was added to a total of 2% (v/v) and incubated for 1 h. at 37°C. Plugs were washed four times (15 min. each time) in Bionano Wash solution, followed by five 15 min. washes in TE Buffer. Finally, ultra-HMW DNA was eluted from the agarose by melting the plugs at 70°C for 2 min. in a thermomixer, allowing the temperature to decrease gradually to 43°C, adding 2 μl agarase and incubated at 43°C for 45 min. The highly viscous DNA samples were further cleaned-up by drop dialysis against TE buffer and quantified using Qubit.

Optical mapping was performed using the Bionano Direct labeling and stain approach (DLS; Bionano Genomics, San Diego, CA) as described in (55). However, only 350 to 500 ng of ultra-HMW DNA was used per reaction. The labeled sample was loaded into a Saphyr G2.3 chip, and molecules separated, imaged, and digitized using a Saphyr analyzer and Compute server.

### Genome size estimation

To estimate the genome size of Ey15-2 from PCR-free reads, we pre-processed the reads and discarded those that aligned to organellar genomes or the Illumina PhiX Control for HiSeq. We trimmed remaining adapters from raw-reads, removed low quality bases and discarded reads shorter than 75 bp (-q 20, 15 --trim-n --minimum-length 75) with cutadapt v2.4 (56). Then, we aligned all reads to the chloroplast and mitochondrial genomes of *A. thaliana* and the bacteriophage phiX174 genome with bwa-mem v0.7.17 (57) to later discard reads that did not aligned to the nuclear genome with a series of Samtools v1.9 (58) commands. To obtain paired-reads alignments in which read1 was unmapped and read2 was mapped, we used ’samtools view -b -f 4 -F 264’. Conversely, to obtain paired-reads alignments in which read1 was mapped and read2 was unmapped, we used ’samtools view -b -f 8 -F 260’. And to retrieve paired-reads in which both reads were unmapped we used ’samtools view -b -f 12 -F 256’. Later, we combined all three previous outputs with ’samtools merge’, discarded supplementary alignments with ’samtools view -b -F 2048’ and converted the BAM file to FASTQ format with bedtools ’bedtools bamtofastq’ v2.27.1 (59). To count k-mers we employed the ’count’ (-C -m 21 -s 5G) and ’histo’ commands from Jellyfish v2.3.0 (60) with a k-mer size of 21. Finally, an R-script from the findGSE tool (41) estimated the genome size to be 143.12 Mb.

### CLR assembly

The CLR subreads BAM file was converted to FASTA format with SAMtools v1.7 (58) and subreads shorter than 10 kb (seq -L 10000) were discarded with seqtk v1.3 (61). This file was used as input for Canu v2.0 (29) for assembly with a maximum input coverage of 200x and an estimated genome size of 140 Mb (canu -pacbio-raw <INPUT-reads> genomeSize=140mb maxInputCoverage=200 correctedErrorRate=0.035 utgOvlErrorRate=0.065 trimReadsCoverage=2 trimReadsOverlap=500). To polish the assembled contigs we aligned a 20% subset of the subreads larger than 10 kb with pbmm2 v1.0.0 (align --preset SUBREAD), and used GCpp v1.9.0 with the Arrow algorithm (PacBio® tools: https://github.com/PacificBiosciences/pbbioconda).

### HiFi reads subsets

q20 High Fidelity (HiFi) reads were generated with the Circular Consensus Sequencing tool from PacBio® ccs v6.0.0 (--min-passes 3 --min-length 10 --max-length 60000 --min-rq 0.99). To study the impact of coverage in different HiFi assemblers, the original ∼133x q20 HiFi dataset was subsetted to 125x, 100x, 75x, 50x and 25x with rasusa v0.3.0 (62) assuming a genome size of 140 Mb. For each coverage subset, five replicates were generated using seed values 3, 19, 23, 54 and 70, resulting in 25 subsets.

To assess the impact of read length in different HiFi assemblers, we trimmed all reads in the original HiFi dataset, which had a median read length of 21.5 kb, with the command ’trimfq’ from seqtk v1.3 (61). By trimming 0, 1, 2, 3, and 4 kb from each end of the reads, we generated subsets with median read lengths of 21.5, 19.5 kb, 17.5 kb, 15.5 kb and 13.5, respectively. Afterwards, reads smaller than 2 kb in the resulting subsets were discarded, and we determined that the coverage in the smallest read subset was slightly above 85x. Finally, all sets were subjected to five replicates of downsampling to 85x with rasusa (62) as explained before, resulting in a total of 25 subsets.

### HiFi assemblies

The original HiFi CCS set, in addition to 25 coverage and 25 read length subsets, were each assembled with HiCanu (30), FALCON (11, 23), Hifiasm (31), Peregrine (32) and IPA (33). Identical commands were used for all different subsets per assembler.

HiCanu was used through Canu v2.0 (29, 30) with a maximum coverage threshold above the read depth of all subsets (-assemble -pacbio-hifi genomeSize=140m maxInputCoverage=200). HiFi FALCON assemblies were run by executing the toolkit (11, 23) distributed with the ’PacBio Assembly Tool Suite’ v0.0.8 (falcon-kit 1.8.1; pypeflow 2.3.0; https://github.com/PacificBiosciences/pb-assembly). An example configuration file with detailed assembly parameters used in this study is provided in the dedicated GitHub for this study. The same input HiFi reads used for assembly were further mapped to the resulting contigs with pbmm2 v1.0.0 (align --preset CCS --sort), and polished with Racon v1.4.10 (63). The assemblies performed with Hifiasm (31) only needed the specification of a parameter for small genomes (-f0) and the disabling of purging of duplicated contigs recommended for inbred genomes (-l0). All Ey15-2 subsets were performed with Hifiasm v0.13-r308, while the Col-0 was assembled with Hifiasm v0.16.1-r375. Peregrine v1.6.3 (32) was run using the following command for all assemblies: ’pg_run.py asm index_nchunk=48 index_nproc=48 ovlp_nchunk=48 ovlp_nproc=48 mapping_nchunk=48 mapping_nproc=48 cns_nchunk=48 cns_nproc=48 sort_nproc=48 --with-consensus --shimmer-r 3 --best_n_ovlp 8’. PacBio’s IPA v1.3.1 (33) was used in cluster mode (dist) and skipping phasin (--no-phase) for inbred genomes. pbmm2 v1.0.0 (align --preset SUBREAD), and used GCpp v1.9.0 with the Arrow algorithm (PacBio® tools: https://github.com/PacificBiosciences/pbbioconda).

### Scaffolding with optical maps

Data visualization, map assembly, and hybrid scaffold construction were performed as per manufacturer’s recommendations using Bionano Access v1.5 and Bionano Solve v3.6 (https://bionanogenomics.com/support/software-downloads). The assembly was performed in pre-assembly mode using parameters ’non-haplotype’ and ’no-CMPR-cut’, without extend-split.

The resulting agp files of the hybrid scaffolds (which can be found here: https://keeper.mpdl.mpg.de/d/216caab287514b1ba2c5/) were manually curated to specifically discard: (1) complete super-scaffolds –and their associated contigs– of organellar DNA, (2) complete super-scaffolds –and their associated contigs– of 45S rDNAs, and (3) isolated contigs “hybridizing” to the 45S rDNA portion of otherwise larger super-scaffolds. A complete list of all super-scaffolds and contigs removed from the Bionano-based scaffolds is provided in Supplementary File 1. Similarly, these contigs were also added to the list of non-scaffolded contigs that was used for the analysis of contig breaks (see below). Edited agp files were converted to fasta format with the script ’ragtag_agp2fasta.py’ from RagTag v1.1.1 (40).

### Reference-based scaffolding

For the evaluation of accuracy and completeness, we scaffolded contigs >150 kb with RagTag v1.1.1 (40) (scaffold -q 60 -f 10000 -I 0.5 --remove-small) using a hard-masked version of TAIR10 as reference genome. For Col-0, the procedure differed slightly: we scaffolded contigs >100 kb with RagTag v2.0.1 (40) (scaffold -q 60 -f 30000 -I 0.5 --remove-small), also using our hard-masked version of TAIR10 as reference. Since we observed that *in silico* scaffolding is subjected to biases due to structural variants segregating between the genome used as reference and the genome of the accession being scaffolded, we took the precaution of masking regions in the TAIR10 reference genome that could lead to misplacement of contigs. To this end, we used the function ’bedtools maskfasta’ v2.27.1 (59) with ranges corresponding to our own annotation of centromeres, telomeres, organellar nuclear insertions and both 5S and 45S rDNAs (see section below). Since our annotation of centromeres is specific to the satellite repeat CEN180, we decided to also mask large portions on the pericentromeric region in TAIR10 (Chr1:14309681-15438174, Chr2:3602469-3728277, Chr3:13586904-13870733, Chr3:14132986-14225247, Chr4:2919189-2981850, Chr4:3024926-3061554, Chr4:3194356-3263238, Chr4:3950509-4061755, Chr5:11184520-11316773, Chr5:11651274-12065554, Chr5:12807214-12870360).

### Assembly metrics

Contiguity, correctness (base-level accuracy) and completeness of the single CLR and all 255 HiFi assemblies were analyzed using identical commands. For contiguity, since the total contig lengths of the different assemblies varied massively (particularly between assemblers), we employed NG50, instead of N50. We defined NG50 as the sequence length of the shortest contig at 50% of the size of the TAIR10 reference genome (119.14 Mb; (38)). Scaffolded length, correctness and completeness metrics were estimated on scaffolded contigs, whether this step was done with Bionano optical maps or reference-based with RagTag. Therefore, depending on the scaffolding method, the exact values for the complete set (133x) differ slightly between Table 1 and Figure 2d-e. To estimate correctness and completeness, we used Merqury v1.1 (36), which compares k-mers in the *de novo* assemblies to those found in the raw PCR-free Illumina short reads. First, two k-mer databases with ’k=18’ were generated per Illumina paired-end read with Meryl v1.3 (64) and afterwards combined (meryl union-sum). Then, Merqury was run for each assembly using these k-mer counts as databases. Finally, genome-wide consensus quality (QV) and completeness scores were collected. We also calculated Benchmarking Universal Single-Copy Orthologs (BUSCO; v3.0.2; ’-l embryophyta_odb10 -m genome -sp arabidopsis’) scores as an additional estimate of completeness (37). Assembly metrics can be found in Supplementary File 2.

### Gap inspection

To create the ’HiFi + CLR’ assembly of Ey15-2, we used the ’patch’ function (-f 10000 --remove-small --join-only) of RagTag v2.0 (40) with the HiFi-HiFiasm contigs as a target and the CLR-Canu contigs as a query. Then, we used pbmm2 v1.3.0 to align the CLR (align --preset SUBREAD --best-n 1 --min-length 500) and HiFi reads (align --preset CCS --best-n 1 --min-length 500) to the new assembly, and IGV v2.6.3 (46) to visualize “patched” loci. To analyze gaps in our HiFi-Hifiasm assembly of Col-0, we aligned the contigs to the Naish *et al.* assembly (18) with minimap2 v2.17 (65) (-ax asm5) and inspected the loci where adjacent contigs break with IGV v2.6.3 (46). We summarized the results of these analyses in Supplementary File 1.

### Annotation and analysis of repetitive elements

We annotated repetitive elements with a custom pipeline in the CLR-Canu assembly, as well as in HiFi-Hifiasm, HiFi-HiCanu, HiFi-FALCON, HiFi-Peregrine and HiFi-IPA assemblies of Ey15-2 that were based on the complete HiFi set. First, we ran RepeatMasker v4.0.9 (66) (-cutoff 200 -nolow -gff -xsmall) using a custom library that included the six CEN180 repeat clusters defined by Maheshwari *et al.* (2017) (67), the three consensus 5S rDNA units from Simon *et al.* (2018) (48), a reference 45S rDNA unit (68), and the telomere motif “CCCTAAA” (x60). Next, with minimap2 v2.16 (65) (-cx asm5), and using the *A. thaliana* mitochondrial and chloroplast genomes (38) as target references, we identified sequences in our assemblies matching to either organellar genome. The gff2 and paf outputs of RepeatMasker and minimap2, respectively, were reformatted to gff3. Separately, transposable elements (TEs) and other repeat regions were annotated with the package Extensive de-novo TE Annotator (EDTA) v1.9.7 (69) (--step all --sensitive 1 --anno 1 --overwrite 1) that includes various TE annotations tools such as LTRharvest, LTR_FINDER, LTR_retriever, TIR-Learner, HelitronScanner, TEsorter (70–77). Finally, to combine all previous annotations, a series of ’merge’ and ’intersect’ commands from bedtools v2.27.1 (59) were used to avoid any overlap between – sometimes – conflictive repetitive elements with the following hierarchy: organellar sequence > rDNAs > TEs.

To contextualize the contribution of these repetitive elements in the assemblies, we counted their cumulative length separately for scaffolded and non-scaffolded contigs as determined from the previous scaffolding with optical maps analysis. For the analysis of contig breaks, only contigs >10 kb were considered, and we determined what repeat was found closer to each contig edge and no more than 2 kb inwards.

For the analysis of centromere and 5S rDNA copy number variation between Ey15-2 and Col-0, we choose the ’HiFi + CLR’ assembly for Ey15-2 and the HiFi-Hifiasm assembly of Col-0. In addition, we downloaded the Col-0 assemblies of Naish *et al.* (18) from https://github.com/schatzlab/Col-CEN/tree/main/v1.2 and of Wang *et al.* (45) from https://ngdc.cncb.ac.cn/gwh/Assembly/21820/show. To estimate the number of 5S rRNA genes before the assembly process of our HiFi samples, we ran RepeatMasker v4.0.9 (66) (-cutoff 200 -nolow -gff -xsmall) directly on q20 HiFi reads using a custom library that included the canonical sequence of rRNA subunits, and counted the number of 5S rRNA gene matches >100 bp (205, 573 and 363, 615 for Ey15-2 and Col-0, respectively). We normalized this number by the genome-wide read depth obtained with samtools (58) (coverage -r Chr3:1-10000000) after aligning the HiFi reads to their own reference with minimap2 v2.17 (-ax asm20), which was 110.352 for Ey15-2 and 121.864 for Col-0.

### Data manipulation and plotting

Most analysis and visualization of our data was done with R v4.0.2 (78) and RStudio v1.3.1073 (79). R packages ’ggplot2’ (80), ’ggh4x’ (81), ’plyr’ (82), ’data.table’ (83) were instrumental for this study. Alignments between assemblies were visualized with AliTV (35) by making use of the MiniTV wrapper (84). Visualization of pericentromeric regions was done with StainedGlass v0.4 (window=5000 mm_f=10000) (47).

## SUPPLEMENTARY FILES

**Supplementary File 1.** Excel file with three sheets: contigs and super-scaffolds manually curated from the hybrid assemblies of Ey15-2; size estimates and information of CLR patches and Bionano gaps in the HiFi-Hifiasm assembly of Ey15-2; size estimates and information of gaps in the HiFi-Hifiasm assembly of Col-0.

**Supplementary File 2.** Metrics of the assemblies based on subsets of data performed by five HiFi assemblers: Hifiasm, HiCanu, FALCON, Peregrine and IPA.

## SUPPLEMENTARY FIGURES

**Supplementary Figure 1.**
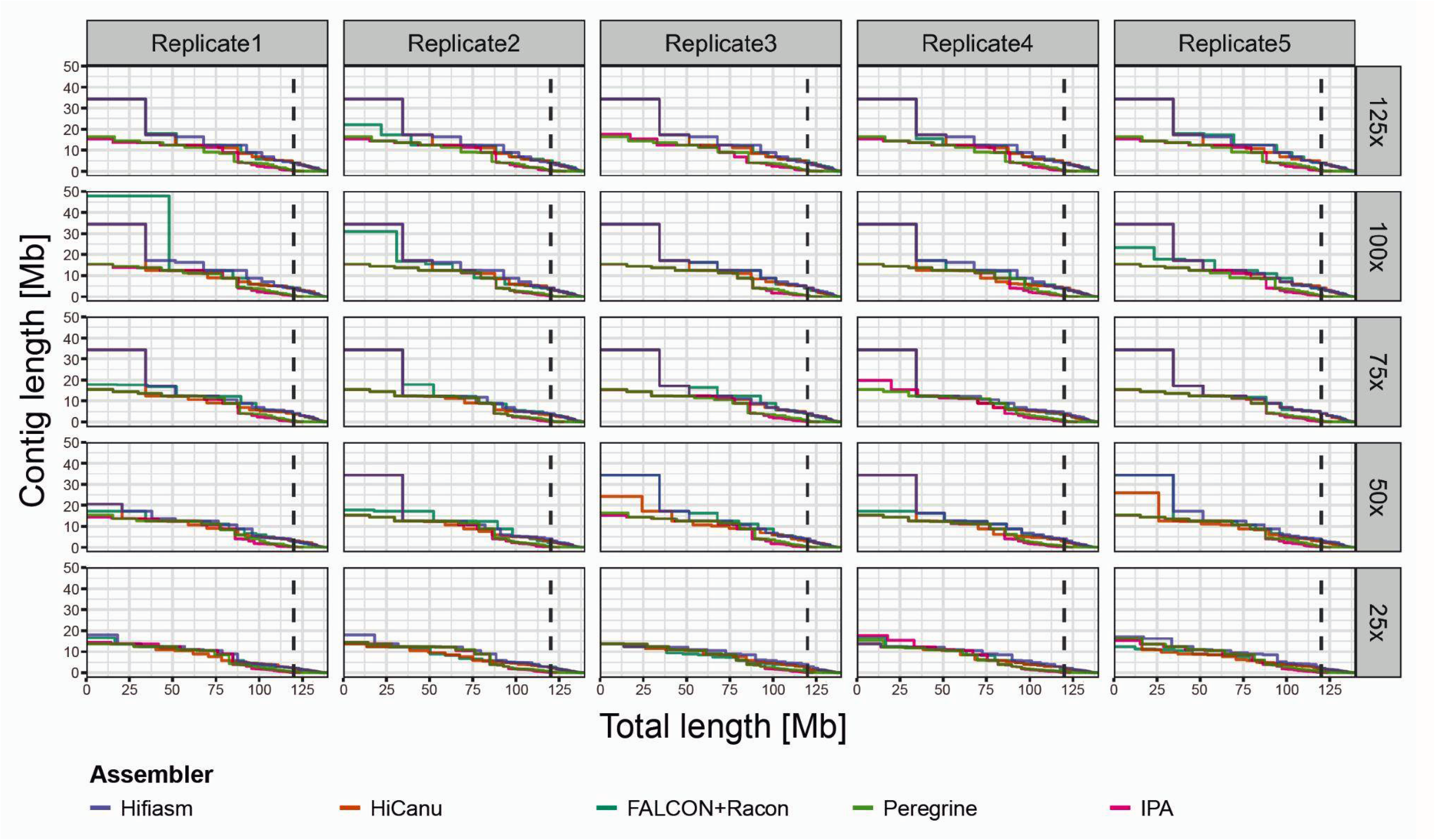
The impact of coverage on five HiFi assemblers. Contiguity plots displaying 125 HiFi assemblies: five assemblers (Hifiasm, HiCanu, FALCON, IPA and Peregrine), five coverage subsets (rows; 125x, 100x, 75x, 50x, 25x), with five replicates each (columns). For each assembly, the cumulative contig length (ordered from largest to shortest) is plotted over the estimated genome size of *A. thaliana* accession Ey15-2 (∼140 Mb). The vertical dashed line indicates the size of the reference genome TAIR10.

**Supplementary Figure 2.**
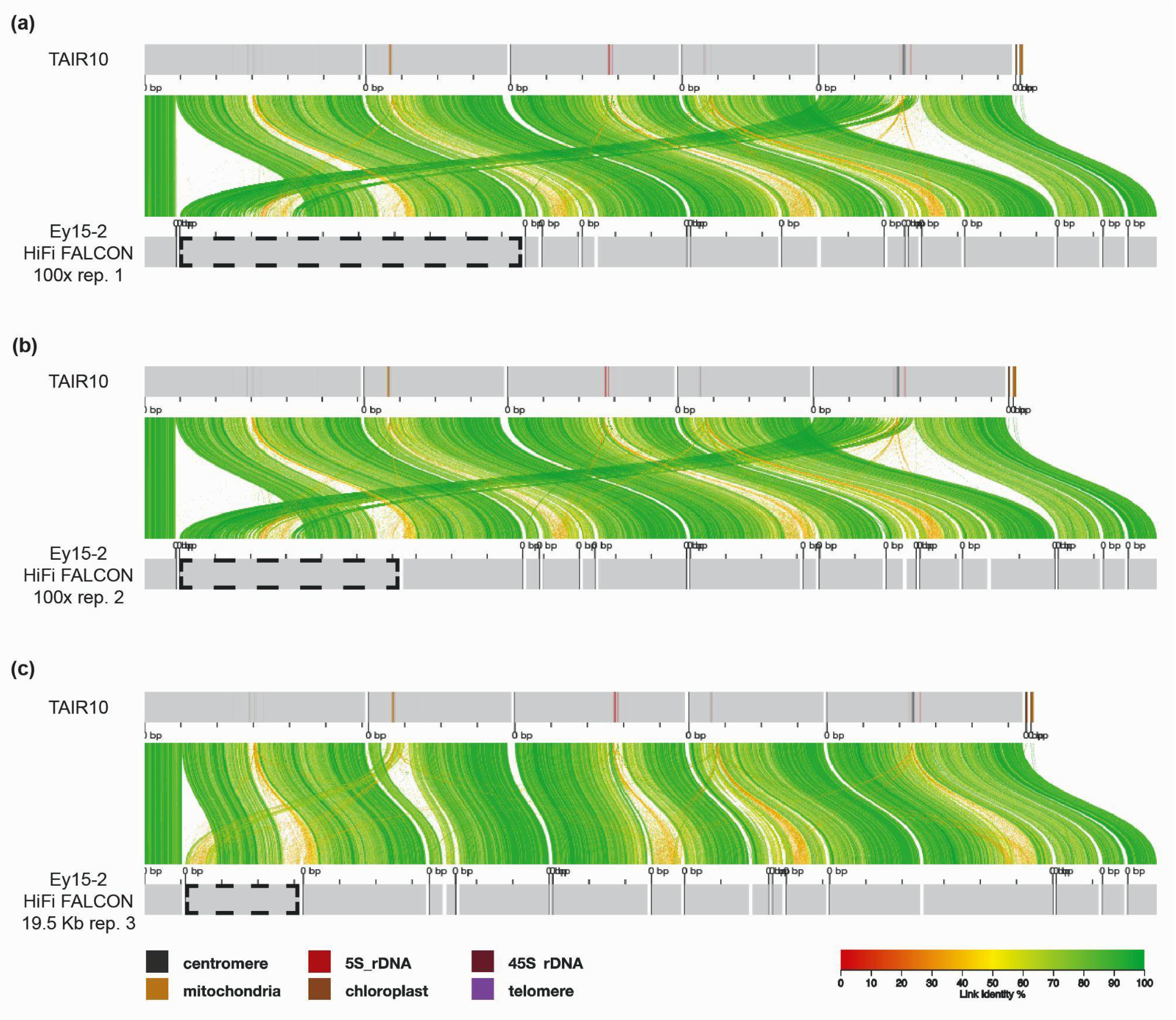
Chimeric contigs produced by the HiFi assembler FALCON. Alignment of the reference genome TAIR10 and the contigs assembled by FALCON (23) with **(a)** coverage subset 100x replicate 1, **(b)** replicate 2, **(c)** and median read length subset 19.5 kb replicate 3 visualized by AliTV (35). Co-linear horizontal gray bars represent chromosomes or contigs, with sequence annotated as repetitive elements (centromeres, 5S and 45S rDNAs, telomeres, mitochondrial and chloroplast nuclear insertions) displayed as shades. Only contigs > 1 Mb are shown. Distance between ticks equals 10 Mb. Colored ribbons connect corresponding regions in the alignment.

**Supplementary Figure 3.**
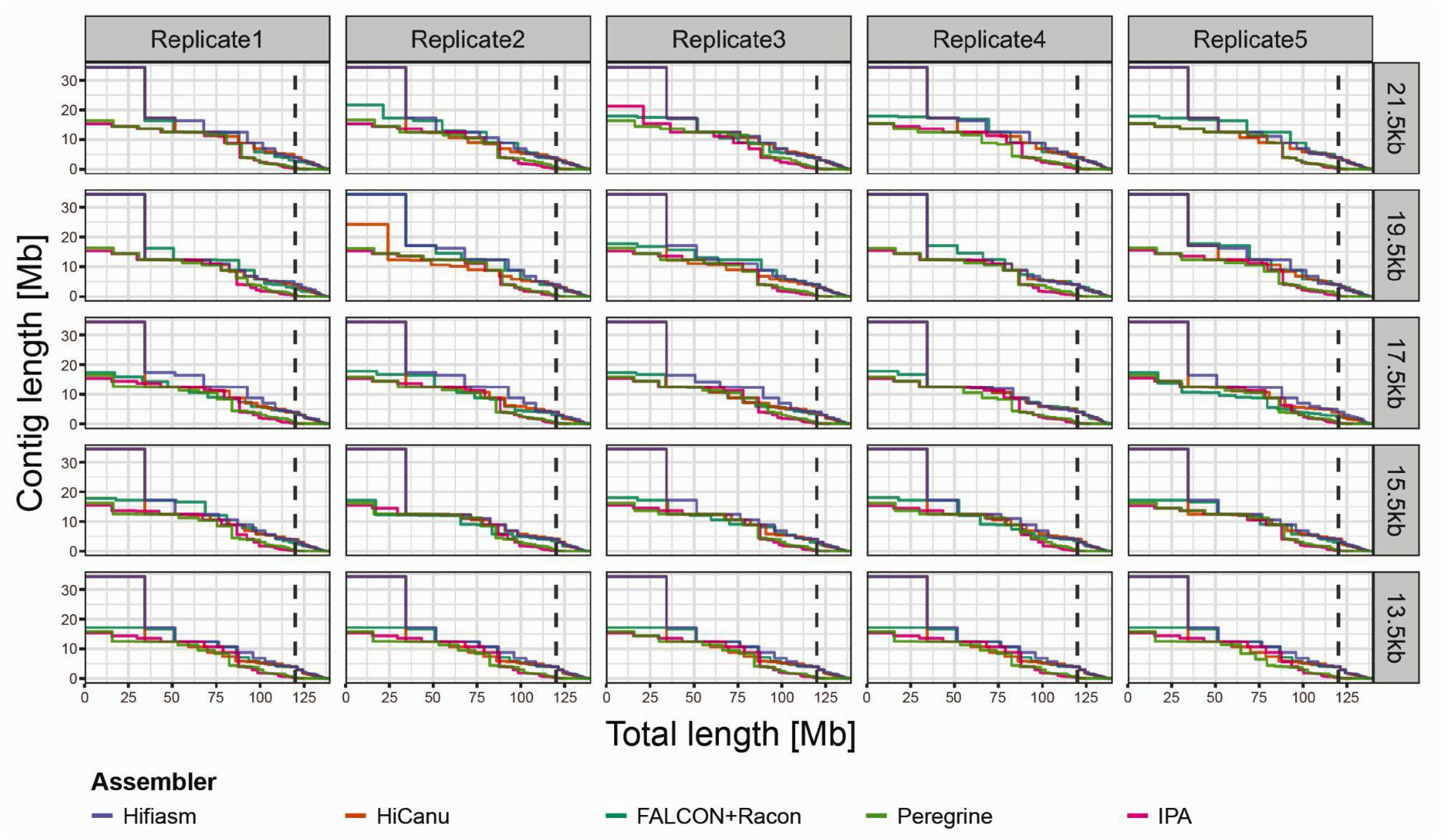
The impact of read length on five HiFi assemblers. Contiguity plots displaying 125 HiFi assemblies: five assemblers (Hifiasm, HiCanu, FALCON, IPA and Peregrine), five median read length subsets (rows; 21.5 kb, 19.5 kb, 17.5 kb, 15.5 kb and 13.5 kb), with five replicates each (columns). For each assembly, the cumulative contig length (ordered from largest to shortest) is plotted over the estimated genome size of *A. thaliana* accession Ey15-2 (∼140 Mb). The vertical dashed line indicates the size of the reference genome TAIR10.

**Supplementary Figure 4.**
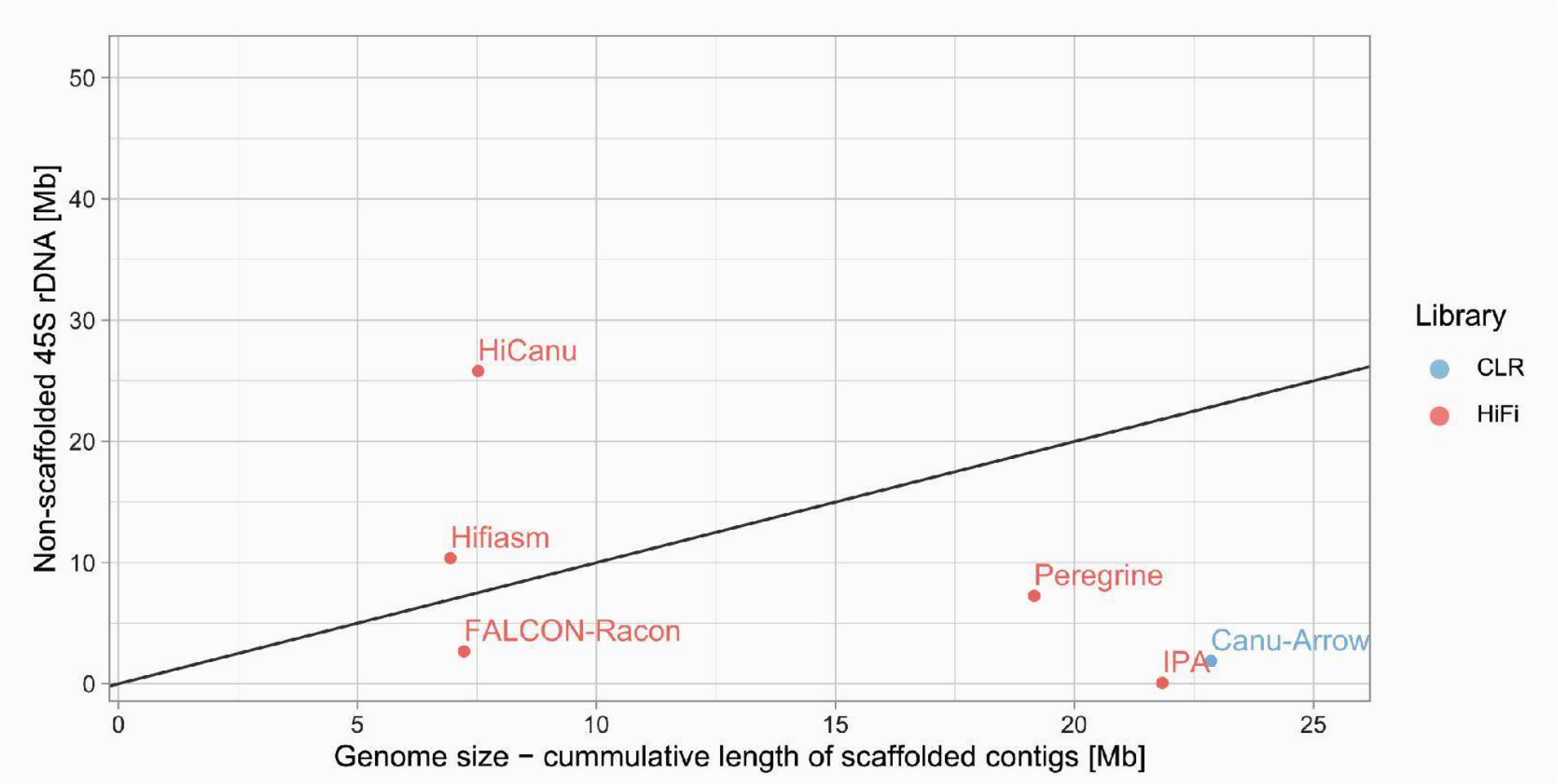
Correlation between the missing portion of the genome and the non-scaffolded 45S rDNA sequence for various assemblers. The solid line indicates the one-to-one relationship between both axes.

**Supplementary Figure 5.**
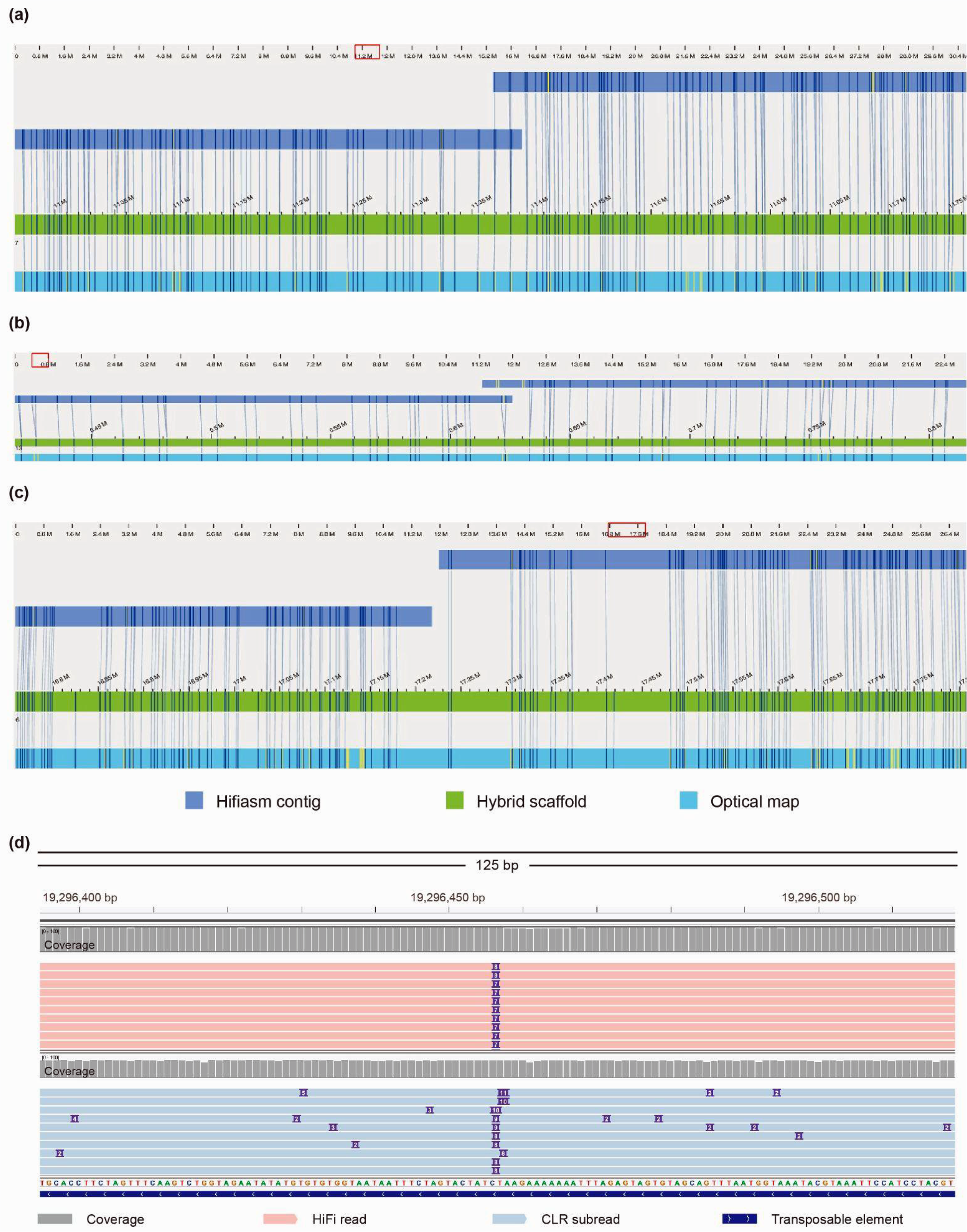
Gaps and overlapping contigs resolved in hybrid scaffolds. **(a)** Hybrid scaffold (in green) between Hifiasm contigs (in dark blue) and the Bionano optical map (in light blue) of Ey15-2 at a locus in Chr5 that the Hifiasm assembly alone was not able to resolve a pair of overlapping contigs. (**b**) Similar to (a) but at a locus in Chr2 that evidenced the inconsistency in labeling pattern for one of the contig edges. **(c)** Similar to (a) but at a locus in Chr3 where adjacent contigs did not overlap, creating a gap. Vertical lines connect consistent labeling positions at DLE-1 recognition sites between long read contigs and Bionano optical maps. **(d)** Visualization with IGV (46) of aligned HiFi reads (in red; top) and CLR (in blue; bottom) over Chr5:19296395-19296519 in the HiFi + CLR assembly. In the original HiFi-Hifiasm scaffold, the corresponding locus had a gap due to an unresolved contig overlap (a), but RagTag identified and fixed it, albeit leaving a 7 bp deletion, evidenced by the alignment of long reads.

**Supplementary Figure 6.**
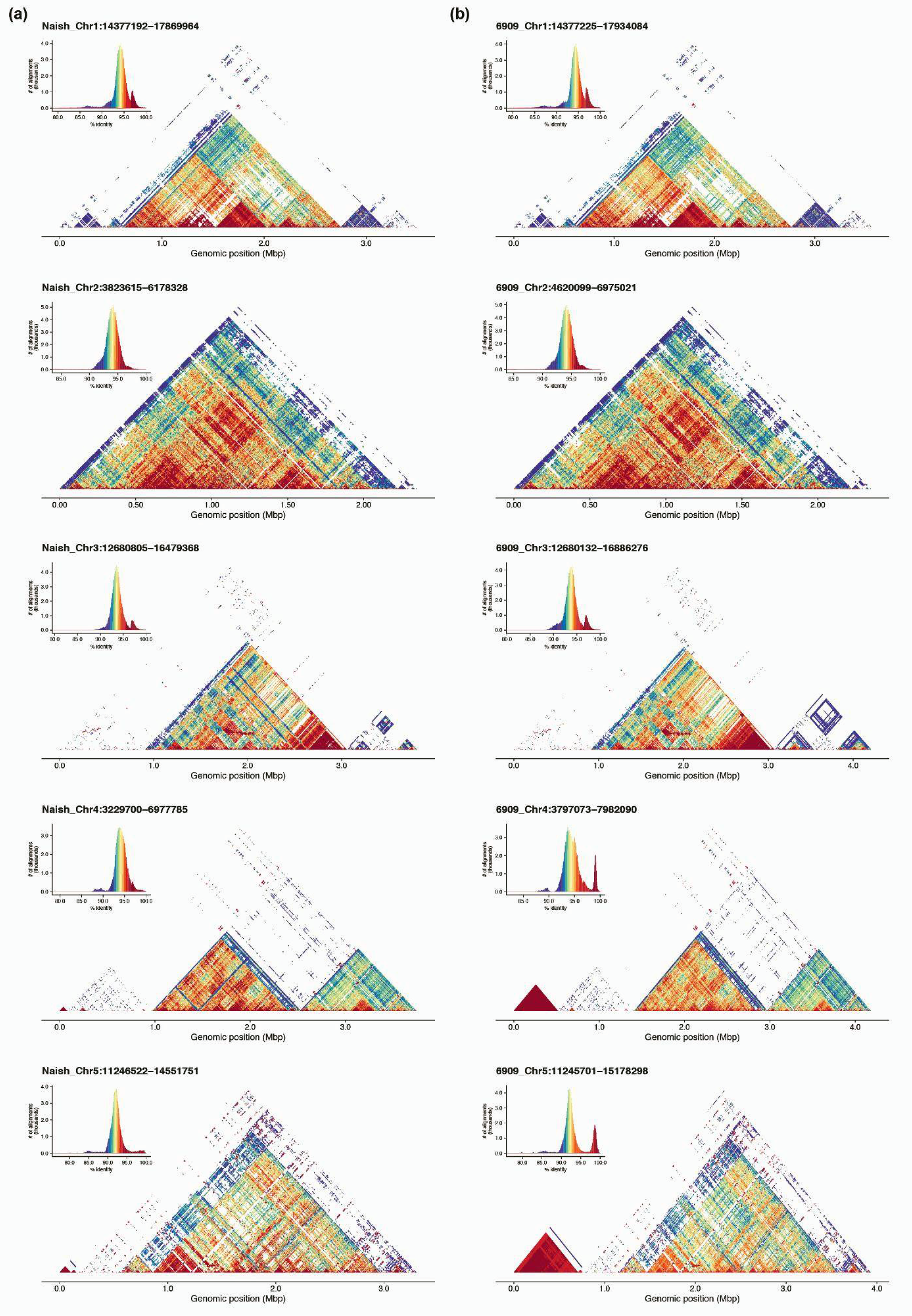
Pericentromeric regions in the Col-0 assemblies from **(a)** Naish *et al.* (18) and **(b)** our HiFi-Hifiasm visualized by StainedGlass (47). While centromeres are largely consistent, 5S rDNA clusters are not. Histograms of the colored percent identity per centromere are also shown.

**Supplementary Figure 7.**
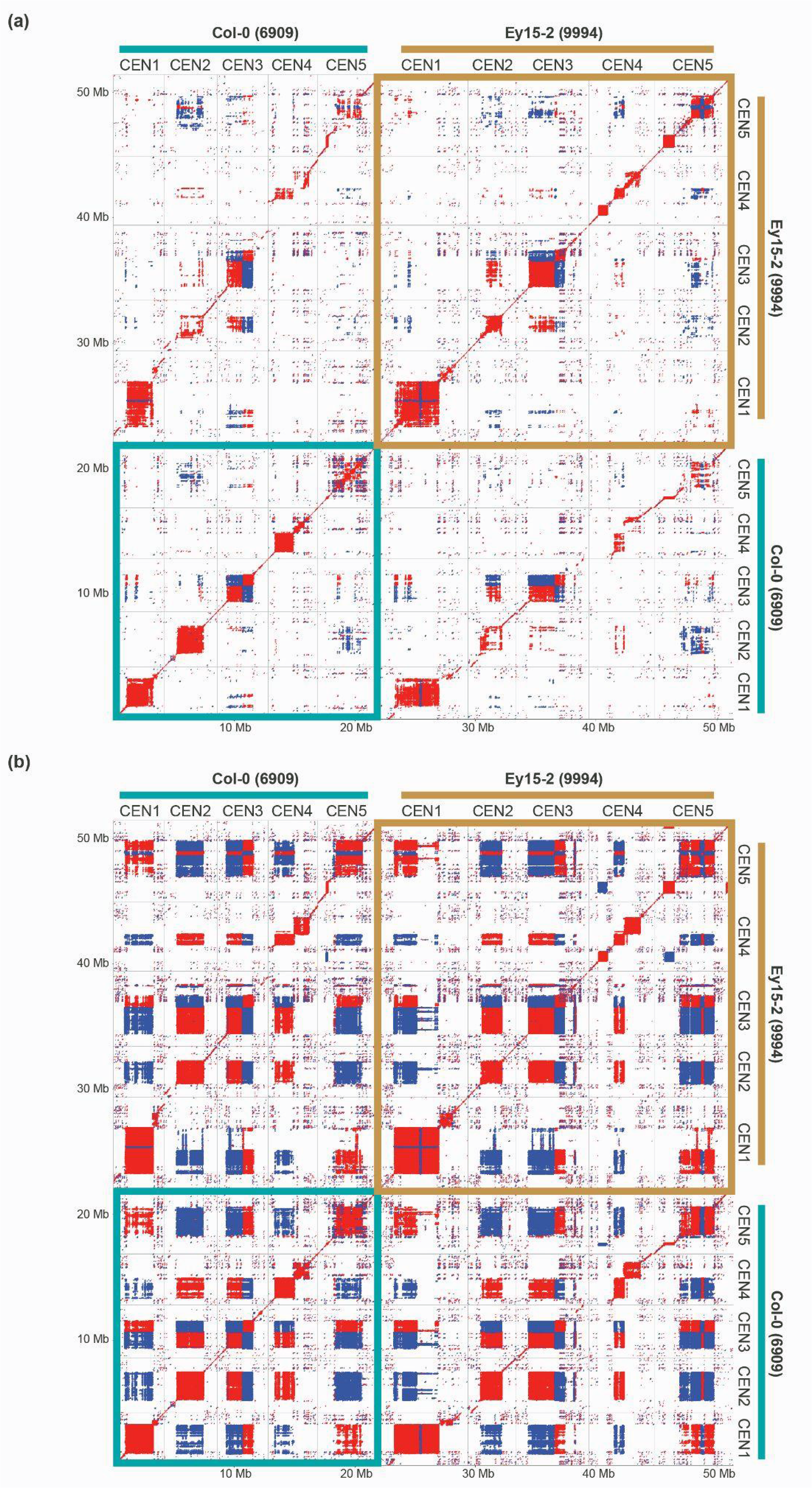
Dotplot analysis comparing the five pericentromeric regions of Col-0 and Ey15-2. **(a)** Using a search window of 178 bp. **(b)** Using a search window of 130 bp. Red and blue shading indicate detection of similarity on the same or opposite strands, respectively.

**Supplementary Figure 8.**
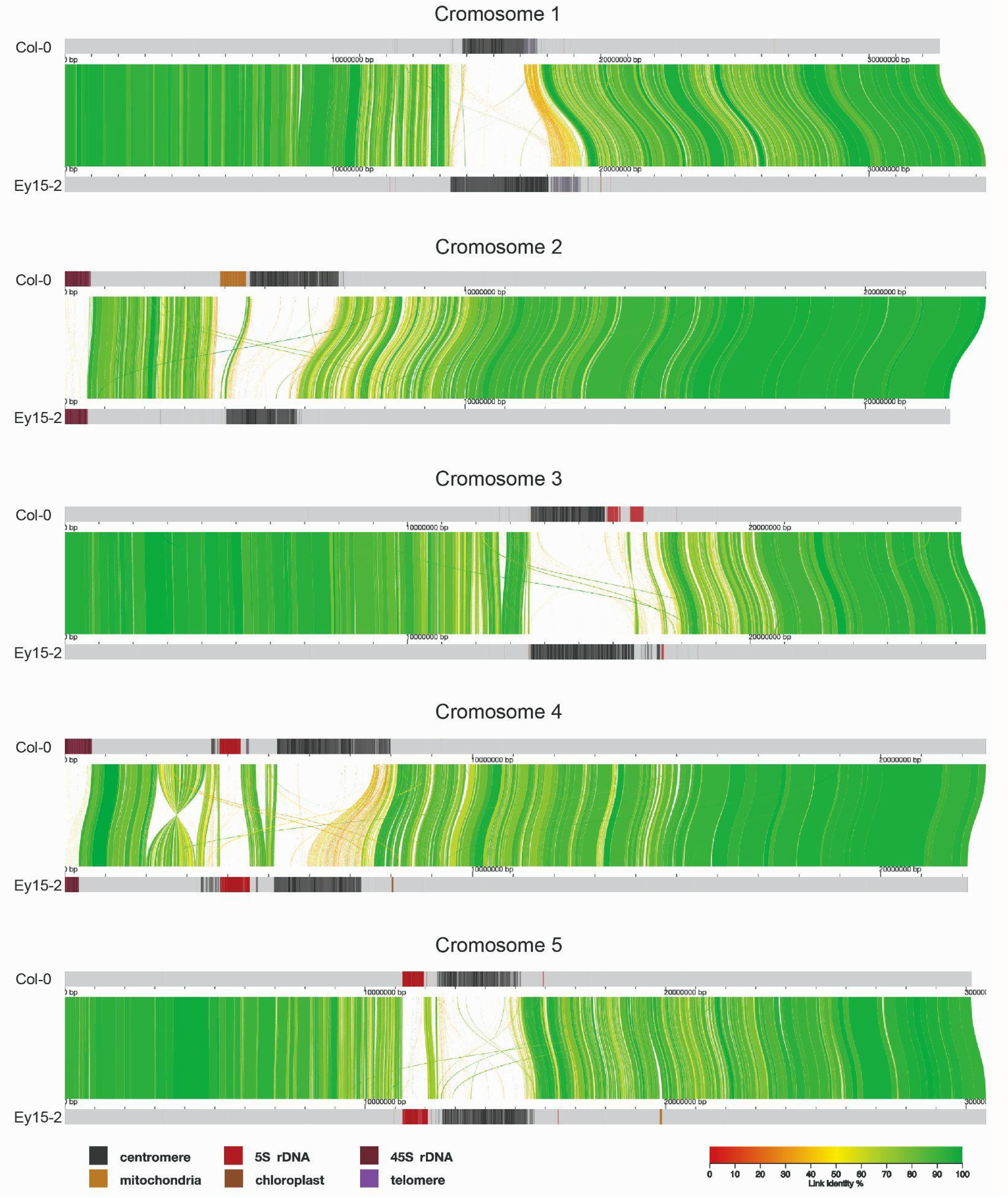
Alignment of the HiFi + RagTag scaffolds of Col-0 and the HiFi + CLR assembly of Ey15-2 visualized by AliTV (35). Co-linear horizontal gray bars represent the five chromosomes in *A. thaliana*, with sequence annotated as repetitive elements (centromeres, 5S and 45S rDNAs, telomeres, mitochondrial and chloroplast nuclear insertions) displayed as shades. Distance between ticks equals 1 Mb. Colored ribbons connect corresponding regions in the alignment.

**Supplementary Figure 9.**
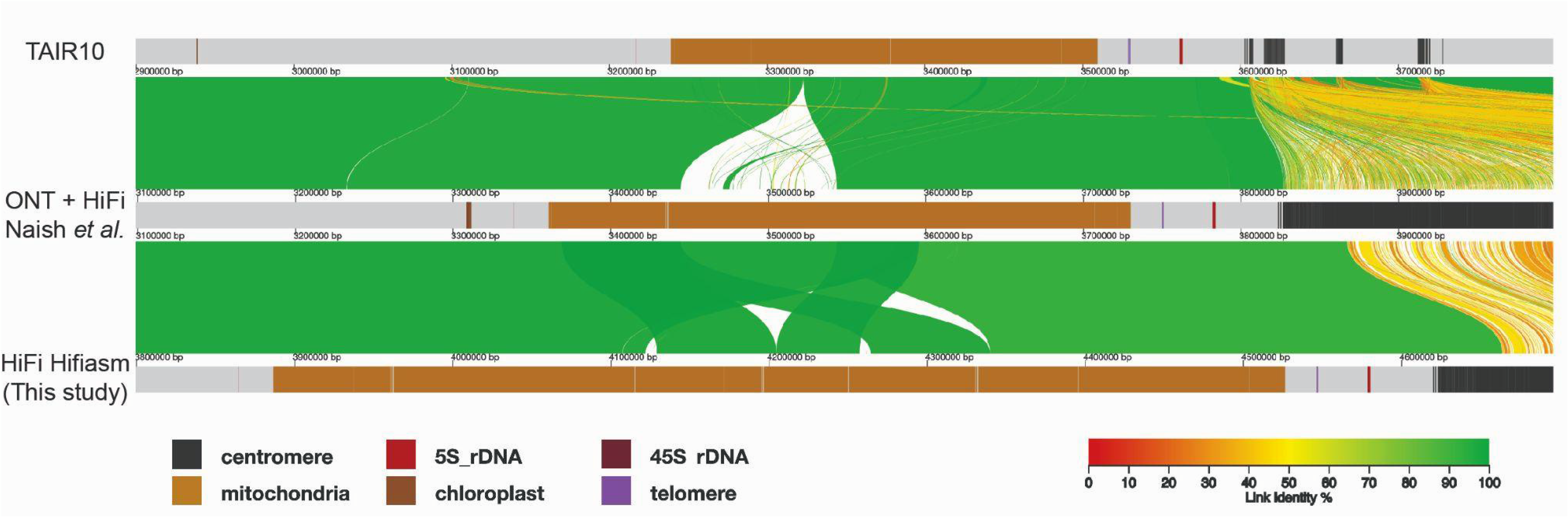
Alignment of multiple assembly versions of the mitochondrial DNA insertion downstream of the centromere in Chromosome 2 in Col-0 visualized by AliTV (35). Co-linear horizontal gray bars represent portions of the TAIR10 reference genome (*top*; (1)), the assembly from Naish et al. (*middle*; (18)) and our HiFi-Hifiasm assembly of Col-0 (*bottom*), with sequence annotated as repetitive elements (centromeres, 5S and 45S rDNAs, telomeres, mitochondrial and chloroplast nuclear insertions) displayed as shades. Distance between ticks equals 100 kb. Colored ribbons connect corresponding regions in the alignment.

## Notes

https://keeper.mpdl.mpg.de/d/216caab287514b1ba2c5/

https://github.com/frabanal/A.thaliana_CLR_vs_HiFi

### Competing Interest Statement

The authors have declared no competing interest.

### Summary of Updates

This version of the manuscript has been revised solely to update the quality of the main figures attached at the end of the composite pdf file.

